# *Botrytis cinerea* detoxifies the sesquiterpenoid phytoalexin rishitin through multiple metabolizing pathways

**DOI:** 10.1101/2024.03.08.584175

**Authors:** Abriel Salaria Bulasag, Akira Ashida, Atsushi Miura, Sreynich Pring, Teruhiko Kuroyanagi, Maurizio Camagna, Aiko Tanaka, Ikuo Sato, Sotaro Chiba, Makoto Ojika, Daigo Takemoto

**Affiliations:** Graduate School of Bioagricultural Sciences, Nagoya University, Chikusa, Nagoya, 464-8601, Japan; College of Arts and Sciences, University of the Philippines Los Baños, College, Laguna 4031, Philippines

## Abstract

*Botrytis cinerea* is a necrotrophic pathogen that infects across a broad range of plant hosts, including high-impact crop species. Its generalist necrotrophic behavior stems from its ability to detoxify structurally diverse phytoalexins. The current study aims to provide evidence of the ability of *B. cinerea* to tolerate the sesquiterpenoid phytoalexin rishitin, which is produced by potato and tomato. While the growth of potato pathogens *Phytophthora infestans* (late blight) and *Alternaria solani* (early blight) was severely inhibited by rishitin, *B. cinerea* was tolerant to rishitin. After incubation of rishitin with the mycelia of *B. cinerea*, it was metabolized to at least six oxidized forms. Structural analysis of these purified rishitin metabolites revealed a variety of oxidative metabolism including hydroxylation at C7 or C12, ketone formation at C5, and dihydroxylation at the 10,11-olefin. Six rishitin metabolites showed reduced toxicity to *P. infestans* and *A. solani*, indicating that *B. cinerea* has at least 5 distinct enzymatic reactions to detoxify rishitin. Four host-specialized phytopathogenic *Botrytis* species, namely *B. elliptica, B. allii, B. squamosa,* and *B. tulipae* also had at least a partial ability to metabolize rishitin as *B. cinerea*, but their metabolic capacity was significantly weaker than that of *B. cinerea*. These results suggest that the ability of *B. cinerea* to rapidly metabolize rishitin through multiple detoxification mechanisms could be critical for its pathogenicity in potato and tomato.

## 1. Introduction

When plant cells recognize an attempt of infection by pathogens, they may produce low molecular weight antimicrobial substances as a countermeasure. These substances are collectively referred to as phytoalexins, and plants have been shown to produce phytoalexins of diverse chemical groups. The production of phytoalexins is a key plant defensive strategy against plant pathogens (Hammerschmidt 1999; Ahuja et al., 2012; Shibata et al., 2016; Imano et al., 2022). Through evolutionary lineages and plant-pathogen contact events, these phytoalexins were sculpted and diversified, giving rise to a vast range of structurally different molecules. These phytoalexins provide sufficient defense against the majority of non-host fungal diseases due to their structural variations, precise timing, and synergistic action (Pedras and Abdoli, 2017; Allan et al., 2019; Newman and Derbyshire, 2020). The ability to detoxify phytoalexins can therefore have a decisive impact on whether a pathogen can successfully infect a given plant. In the case of pathogens with wide host ranges, such as *Sclerotinia* and *Botrytis*, the detoxification of diverse phytoalexins appears to have been a prerequisite for their host range expansion (Westrick et al., 2021; Kuroyanagi et al., 2022, Kusch et al., 2022, Bulasag et al., 2023).

Among the best-known phytoalexin metabolizing enzymes is pisatin demethyltransferase (PDA) produced by the pea pathogen *Nectria haematococca* mating population VI. Knockout of the PDA gene impaired the virulence of *N. haematococca* on peas, clearly demonstrating directly the importance of PDA for successful infection by this pathogen. A correlation between pathogenicity to pisatin-producing plants and the ability to produce active PDA has been reported, indicating that it is the enzyme that determines the host range of *N. haematococca* and *Fusarium oxysporum* races (Wasmann and VanEtten, 1996; Coleman et al., 2011).

*B. cinerea* is a necrotrophic plant pathogen notorious for its ability to infect hundreds of plant species, many of which produce phytoalexins. It follows, therefore, that *B. cinerea* must excel at coping with a diverse range of phytoalexins. Previously, we have reported that *B. cinerea* can metabolize the sesquiterpenoid phytoalexin capsidiol, a major phytoalexin of *Nicotiana* (e.g. tobacco) and *Capsicum* (e.g. chill pepper) species, into non-toxic capsenone using the dehydrogenase BcCPDH (Kuroyanagi et al., 2022). In contrast, rishitin, a structurally similar phytoalexin found in certain *Solanum* species, was not targeted by BcCPDH. This study aims to profile the enzymatic metabolization of rishitin by *B. cinerea* and to determine the toxicity of these products.

## 2. Materials and methods

### 2.1. Biological material and growth conditions

*Botrytis cinerea* strain AI18 was isolated from strawberry (Kuroyanagi et al., 2022). The following pathogen stocks were obtained from the stock center of the Ministry of Agriculture, Forestry and Fisheries (MAFF), Japan: *Alternaria solani* strain KL1 isolated from potato (MAFF244036), *A. brassicicola* strain BA31 isolated from Broccoli (MAFF242993), *B. elliptica* strain S0210 isolated from *Lilium* sp. (MAFF306626), *B. allii* strain Yuki11-1 isolated from onion (MAFF307143), *B. squamosa* strain 5ND4 isolated from Chinese chive (MAFF244973) and *B. tulipae* strain 4-3 isolated from tulip (MAFF245230). The stocks were grown on potato dextrose agar (PDA) at 23°C. *Phytophthora infestans* strain 08YD1 (Shibata et al., 2010) was grown on rye media at 20°C.

### 2.2. Incubation of pathogens in phytoalexin solutions and detection of rishitin and its metabolites using LC/MS

Synthesized rishitin (Murai et al., 1975) was provided by former Prof. Akira Murai (Hokkaido University, Japan). For the incubation in rishitin or its metabolites, mycelia plug (approx.1 mm^3^) were excised from the growing edge of the colony using a dissection microscope (Stemi DV4 Stereo Microscope, Carl Zeiss, Oberkochen, Germany) and submerged in 50 μl of water or rishitin solution in a sealed 96 well clear plate. The plate was incubated at 23°C for the indicated time. Outgrowth of hyphae was monitored under a BX51 light microscope (Olympus, Tokyo, Japan) and measured using ImageJ software (Schneider et al., 2012).

For LC/MS measurement, the supernatant of mycelial blocks (1mm^3^) incubated for 72 h in 50 µl of 100 µM rishitin in a sealed 96-well plate well was mixed with 50 µl acetonitrile and measured by LC/MS (Accurate-Mass Q-TOF LC/MS 6520, Agilent Technologies, Santa Clara, CA, USA) with ODS column Cadenza CD-C18, 75 x 2 mm ODS column (Imtakt, Kyoto, Japan), using a 10-100% MeCN (20 min), 0.2 ml/min.

### 2.3 Separation and structural determination of rishitin metabolites

#### 2.3.1. General

UV spectra and specific rotations were obtained by a V-730 BIO spectrophotometer and a D-1010 polarimeter (Jasco, Tokyo, Japan), respectively. NMR spectra were investigated on an Avance ARX400 spectrometer (Bruker Bio Spin, Yokohama, Japan). The chemical shifts (ppm) were referenced to the solvent residual peak at δ_H_ 7.26 ppm (CDCl_3_). HPLC purification was performed by using a high-pressure gradient system equipped with PU-2087plus pumps, a DG-2080-53 degasser, and a UV-2075plus detector (Jasco). LC/MS was measured by a 6520 Accurate-Mass Q-TOF spectrometer (Agilent Technologies) connected to an 1100 high-performance liquid chromatography (HPLC) system (Agilent).

#### 2.3.2. Separation of rishitin metabolites

The supernatants obtained from two 10 ml and two 30 ml of 500 µM rishitin solution in CM media incubated with *B. cinerea* mycelia for over 5 days were combined and extracted twice with EtOAc (60 ml). The organic layers were combined and concentrated, and the obtained residual oily material was separated by HPLC [Develosil ODS-UG-5 (10 mm i.d. × 250 mm) (Nomura Chemical, Seto, Aichi, Japan), 15-55% MeCN (40 min), 3 ml/min, detected at 205 nm], yielding **1a** (0.11 mg, *t*_R_ = 9.9 min), **1b** (0.11 mg, *t*_R_ = 10.4 min), **2** (0.28 mg, *t*_R_ = 17.2 min), **3** (0.23 mg, *t*_R_ = 19.3 min), **4** (0.09 mg, *t*_R_ = 24.9 min), and the fraction containing **5** (*t*_R_ = 25.8 min). The **5**-containing fraction was further purified by HPLC [Develosil ODS-UG-5 (10 mm i.d. × 250 mm), 50-90% MeOH (40 min), 2.8 mL/min, detected at 210 nm], giving pure **5** (0.16 mg, *t*_R_ = 21.5 min) (Supplementary Fig. 1). These metabolites were subjected to high-resolution MS and NMR analyses.

**1a**: [α]^20^_D_ +37 (*c* 0.01, CHCl_3_); high-resolution mass spectra (HR MS) *m/z* 279.1554 (calcd for C_14_H_24_O_4_Na [M+Na]^+^ 279.1567), *m/z* 221.1519 (calcd for C_14_H_21_O_2_ [M-OH-H_2_O]^+^ 221.1536), *m/z* 203.1417 (calcd for C_14_H_19_O [M-OH-2H_2_O]^+^ 203.1430). **1b**: [α]^20^_D_ +57 (*c* 0.01, CHCl_3_); HR MS *m/z* 279.1548 (calcd for C_14_H_24_O_4_Na [M+Na]^+^ 279.1567), *m/z* 221.1521 (calcd for C_14_H_21_O_2_ [M-OH-H_2_O]^+^ 221.1536), *m/z* 203.1416 (calcd for C_14_H_19_O [M-OH-2H_2_O]^+^ 203.1430). **2**: [α]^20^_D_ -120 (*c* 0.026, CHCl_3_); HR MS *m/z* 261.1440 (calcd for C_14_H_22_O_3_Na [M+Na]^+^ 261.1461), *m/z* 221.1527 (calcd for C_14_H_21_O_2_ [M-OH]^+^ 221.1536), *m/z* 203.1419 (calcd for C_14_H_19_O [M-OH-H_2_O]^+^ 203.1430). **3**: [α]^20^_D_ -39 (*c* 0.024, CHCl_3_); HR MS *m/z* 261.1445 (calcd for C_14_H_22_O_3_Na [M+Na]^+^ 261.1461), *m/z* 203.1418 (calcd for C_14_H_19_O [M-OH-H_2_O]^+^ 203.1430). **4**: [α]^20^_D_ -90 (*c* 0.008, CHCl_3_), UV (MeCN) 244 nm (ε 10,700); HR MS *m/z* 259.1294 (calcd for C_14_H_20_O_3_Na [M+Na]^+^ 259.1305), *m/z* 237.1465 (calcd for C_14_H_21_O_3_ [M+H]^+^ 237.1485), *m/z* 219.1359 (calcd for C_14_H_19_O_2_ [M-OH]^+^ 219.1380). **5**: [α]^20^_D_ -94 (*c* 0.014, CHCl_3_); HR MS *m/z* 261.1445 (calcd for C_14_H_22_O_3_Na [M+Na]^+^ 261.1461, *m/z* 203.1412 (calcd for C_14_H_19_O [M-OH-H_2_O]^+^ 203.1430]. ^1^H NMR data are summarized in Table 1.

**Table 1.** ^1^H NMR data for rishitin metabolites (400 MHz, in CDCl_3_).

| Pos | 1a | 1b | 2 | 3 | 4 | 5 |
| --- | --- | --- | --- | --- | --- | --- |
| 1 | 2.09 (m) | 2.12 (m) | 2.07 (m) | 2.08 (m) | 2.35 (m) | 1.35 (m) |
| 2 | 3.23 (brt, 9.0) | 3.24 (t, 9.0) | 3.24 (t, 9.0) | 3.23 (t, 9.2) | 3.32 (t, 9.2) | 3.51 (dt, 3.6, 9.6) |
| 3 | 3.69 (m) | 3.69 (m) | 3.63 (dt, 6.0, 9.0) | 3.66 (m) | 3.6 (dt, 6.4, 9.2) | 3.42 (m) |
| 4 | 2.11 (m),<br>2.22 (m) | 2.11 (m),<br>2.18 (m) | 2.08 (m),<br>2.17 (m) | 2.08 (m),<br>2.19 (m) | 1.99 (m),<br>2.95 (dd, 16.2, 5.8) | 2.44 (d, 7.2) |
| 5 | 1.91 (m),<br>2.1 (m) | 1.88 (m),<br>2.11 (m) | 1.81 (m),<br>2.12 (m) | 1.85 (m),<br>2.02 (m) | - | 5.66 (brd, 5.6) |
| 6 | 1.29 (dq, 5.4, 12.6),<br>1.88 (m) | 1.26 (dq, 5.4, 11.8),<br>1.70 (m) | 1.64 (m),<br>1.88 (m) | 1.58 (m),<br>1.73 (m) | 2.41 (dd, 16.2, 10.2),<br>2.58 (dd, 16.2, 4.4) | 1.88 (dd, 15.6, 11.2),<br>2.14 (m) |
| 7 | 1.74 (m) | 1.77 (m) | - | 2.37 (m) | 2.69 (m) | 2.24 (m) |
| 8 | 1.54 (m),<br>2.14 (m) | 1.68 (m),<br>2.25 (m) | 2.19 (m) | 1.77 (m),<br>2.30 (m) | 2.20 (dd, 18.0, 8.4),<br>2.61 (brd, 18.0) | 1.32 (t, 13.4),<br>2.04 (m) |
| 9 | 1.15 (d, 6.8) | 1.17 (d, 6.4) | 1.17 (d, 6.8) | 1.16 (d, 6.8) | 1.32 (d, 6.8) | 1.15 (d, 6.8) |
| 11 | 3.45 (dd, 10.9, 5.2),<br>3.60 (dd, 10.9, 4.6) | 3.47 (dd, 10.4, 4.8),<br>3.62 (dd, 10.4, 4.0) | 4.84 (s),<br>4.86 (d, 1.2) | 4.84 (s),<br>5.08 (s) | 4.70 (s),<br>4.82 (s) | 4.74 (s),<br>4.77 (s) |
| 12 | 1.12 (s) | 1.13 (s) | 1.81 (s) | 4.14 (s) | 1.74 (s) | 1.75 (s) |
| 2-OH | 2.28 (brd, 3.6) | 2.27 (m) | 2.3 (br) | 2.27 (brs) | 2.6 (brs) | 2.13 (brd, 3.6) |
| 3-OH | 2.16 (brs) | 2.18 (m) | 2.19 (m) | 2.17 (brs) | 2.05 (brs) | 2.21 (brd, 3.6) |
| Other OH | 1.51 (s)<sup>b</sup>,<br>1.81 (m)<sup>c</sup> | 1.51 (s)<sup>b</sup>,<br>1.79 (m)<sup>c</sup> | - | 1.34 (brs)<sup>d</sup> | - | 1.51 (s)<sup>e</sup> |

<sup>a</sup>Multiplicity and coupling constants (in Hz) are in parentheses. <sup>b-e</sup>Shifts for 10-OH, 11-OH, 12-OH, and 8a-OH, respectively.

### 2.5 Determination of antimicrobial activity of rishitin metabolites

Mycelia blocks (approx. 1 mm^3^) of fungal pathogens grown on PDA or oomycete pathogen grown on rye media were excised from the growing edge of the colony using a dissection microscope (Stemi DV4 Stereo Microscope) and submerged in 50 µl of 500 µM rishitin solution or purified rishitin metabolites in a sealed 96 well clear plate. The plate was incubated at 23°C for the indicated time and the outgrowth of hyphae was monitored under a BX51 light microscope (Olympus) and measured using ImageJ software (Schneider et al., 2012).

## 3. Results

### 3.1 Rishitin can inhibit the growth of potato pathogens, Phytophthora infestans and Alternaria solani, but not polyphagous B. cinerea

Rishitin is the major phytoalexin produced in potato tubers and tomato fruits (Katsui et al., 1968; de Wit and Flach, 1979). The sensitivity of potato late blight pathogen, *P. infestans* (oomycete), and potato early blight pathogen, *A. solani* (fungus), to rishitin was tested. The mycelial block of pathogens grown on solid culture media was incubated in 500 µM rishitin for 12 or 24 h and outgrowth of pathogen hyphae was measured. For both pathogens, hyphal growth was significantly inhibited by rishitin (93% for *P. infestans* and 73% for *A. solani*, Fig. 1A and B). An inhibitory effect was also detected for *A. brassicicola* (88 %), which causes black spot disease on Brassicaceae plants such as cabbage, canola, and Arabidopsis (Maude and Humpherson-Jones, 1980; van Wees et al, 2003) (Fig. 1C). In contrast, *Botrytis cinerea* was relatively resistant to rishitin, showing a mere growth inhibition of approximately 20% (Fig. 1D).

**Fig. 1.**
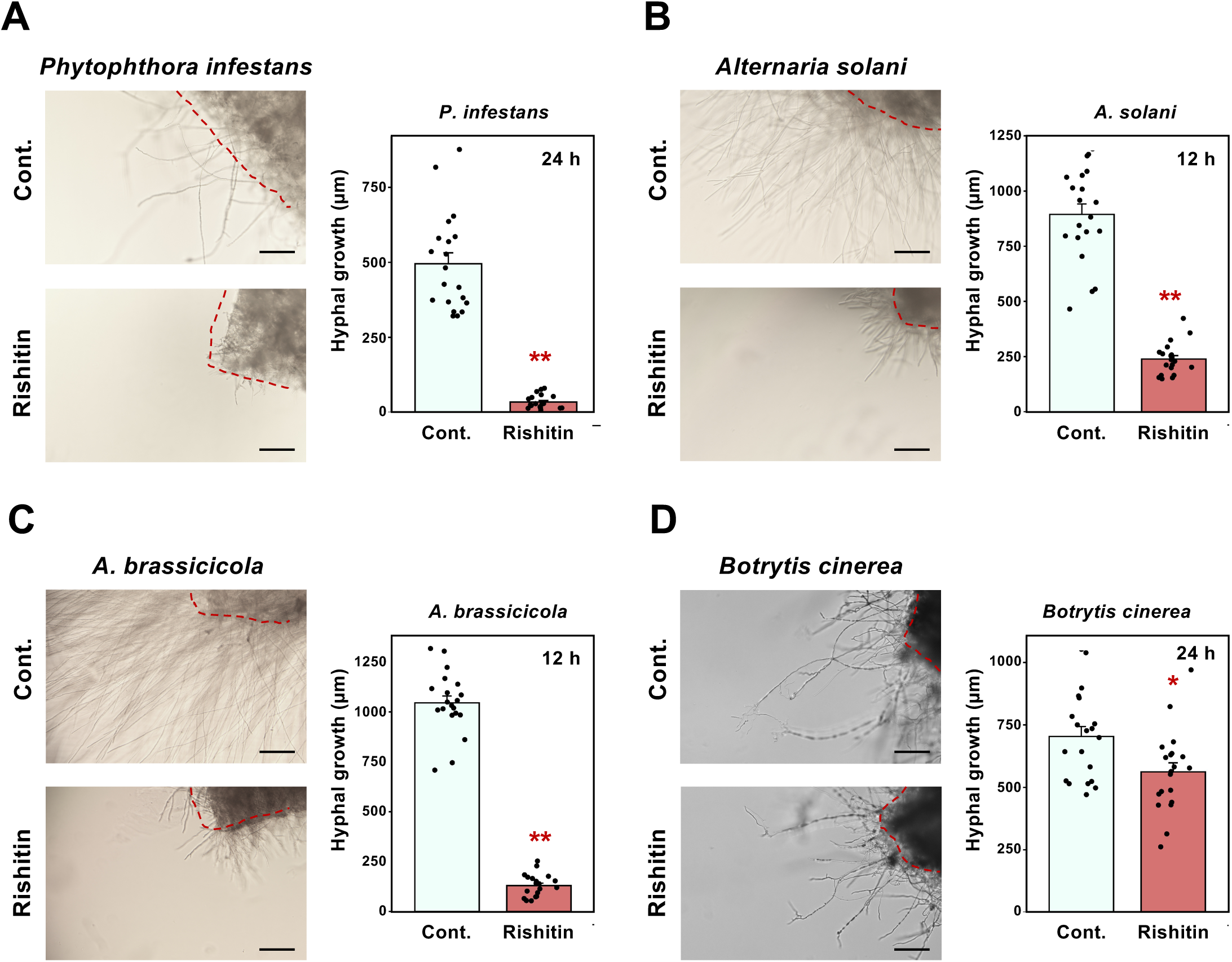
Growth inhibition of phytopathogenic oomycete and fungi by rishitin, a major anti-microbial compound (phytoalexin) produced by potato and tomato. Mycelial blocks (approx. 1 mm^3^) of *Phytophthora infestans* (**A**), *Alternaria solani* (**B**), *A. brassicicola* (**C**), or *Botrytis cinerea* (**D**) were incubated in 50 µl of 0.1% DMSO (Cont.) or 500 µM rishitin. Outgrowth of hyphae from the mycelial block (outlined by dotted red lines) was measured after 12 or 24 h incubation. Bars = 200 µm. Data are means ± SE (n = 20). Data marked with asterisks are significantly different from the control as assessed by the two-tailed Student’s *t*-test: **P < 0.01, *P < 0.05.

### 3.2. Rishitin was metabolized into at least 4 oxidized and 2 oxidized/hydrated forms by B. cinerea

Our previous study indicated that *B. cinerea* can metabolize rishitin into several oxidized forms (Kuroyanagi et al., 2022). To identify these rishitin metabolites, rishitin was incubated with mycelia of *B. cinerea* for 72 h and the metabolites obtained were detected by LC/MS. After the incubation, original rishitin was not detected anymore, and at least 4 oxidized metabolites (m/z = 261.15) were detected (Fig. 2). Moreover, 2 peaks for metabolized rishitin (oxidized followed by hydrated, m/z=279.16) were also detected (Fig. 3), indicating that *B. cinerea* possesses multiple mechanisms to transform rishitin, into presumably less toxic compounds.

**Fig. 2.**
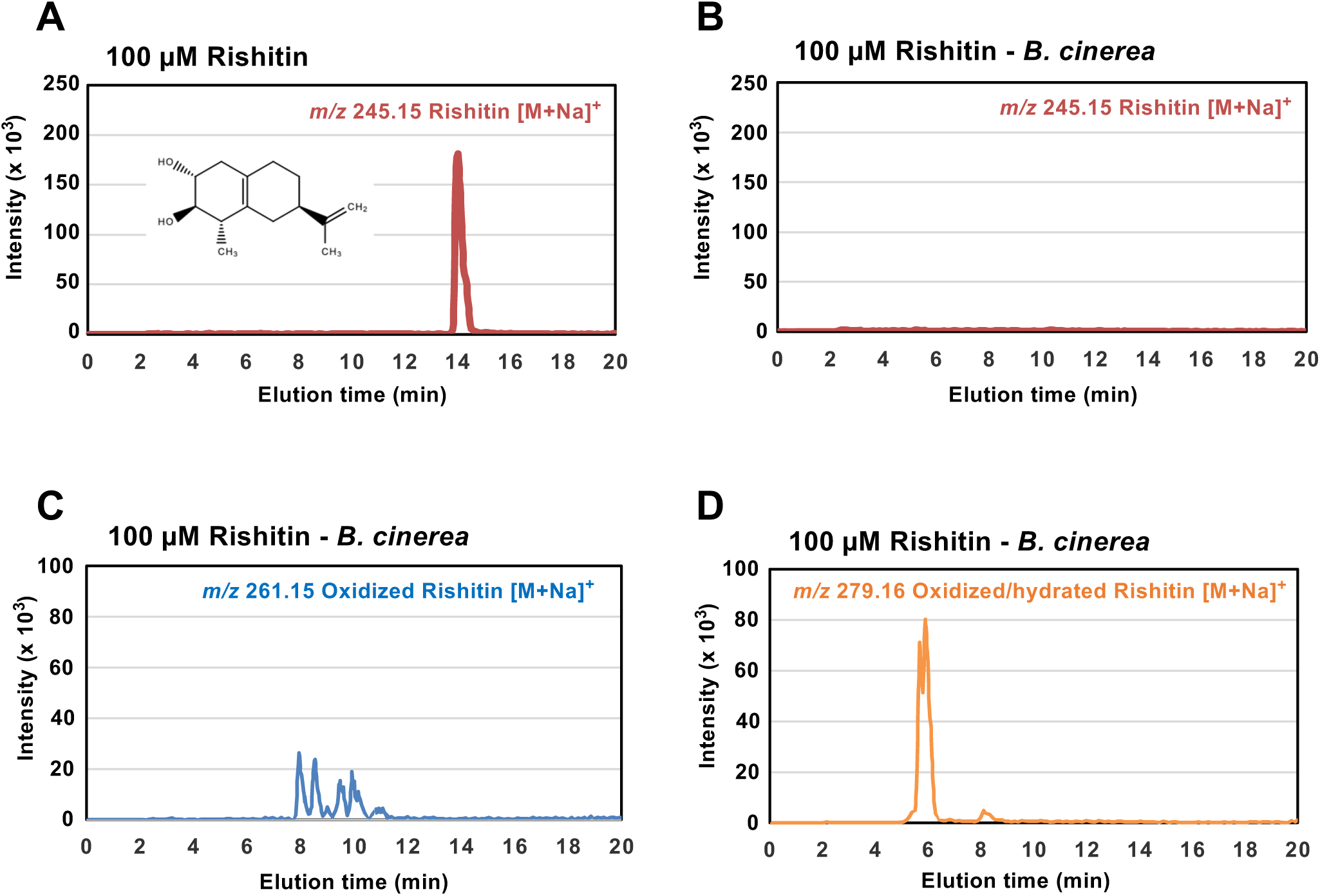
Metabolism of rishitin by *Botrytis cinerea*. Mycelial block (approx. 1 mm^3^) of *B. cinerea* was incubated in 100 µM rishitin for 72 h. Rishitin or its metabolites were detected by LC/MS. (**A, B**) Detection of rishitin (m/z 245.15) incubated without (A) or with (B) *B. cinerea*. (**C, D**) Detection of predicted oxidized rishitins (C, m/z 261.15) and oxidized and hydrated rishitins (D, m/z 279.16) after incubation with *B. cinerea*.

**Fig. 3.**
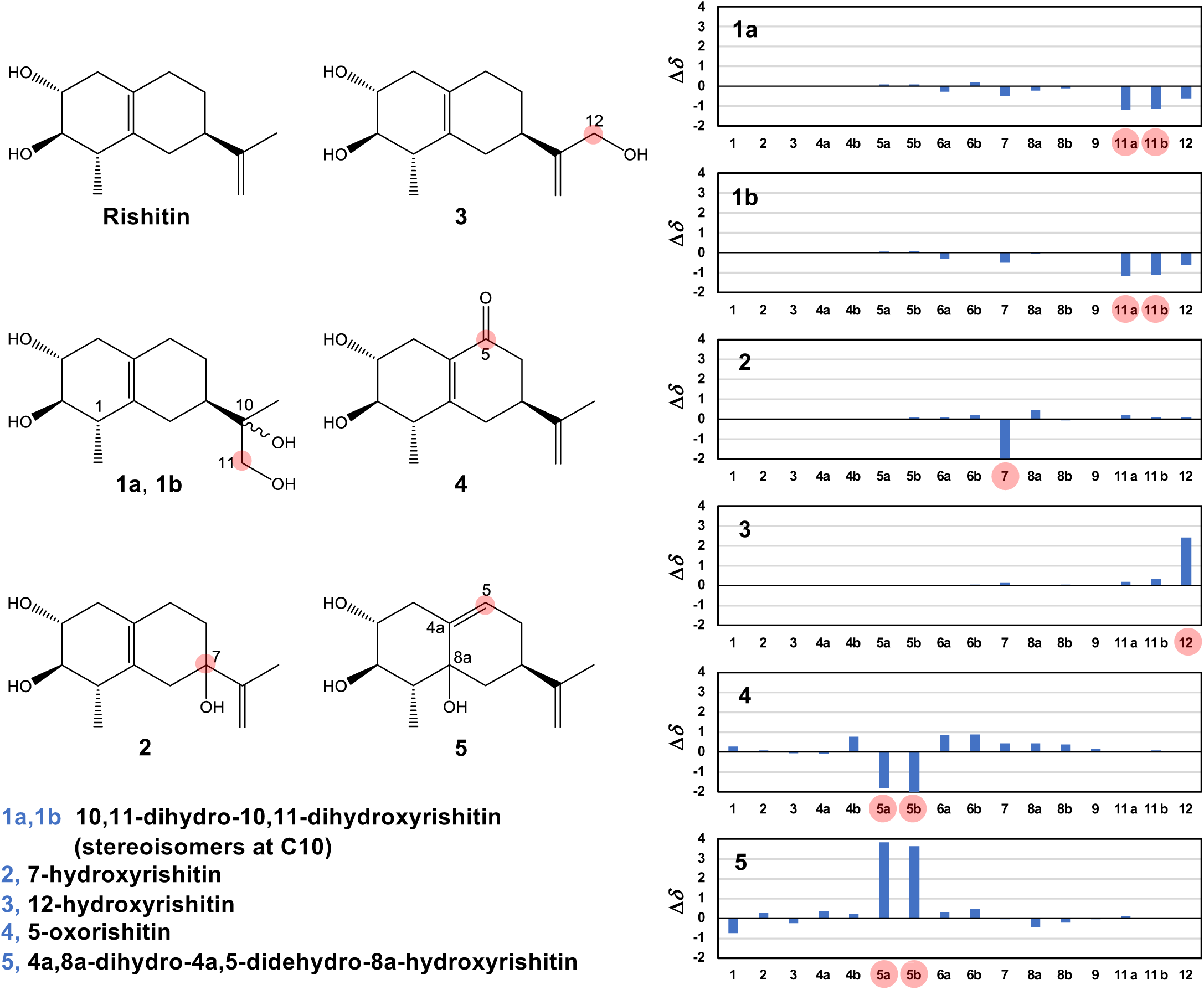
(left) Chemical structure of rishitin metabolites produced by *Botrytis cinerea*. The position numbering of the ring system follows the IUPAC nomenclature. (right) Proton chemical shift differences between metabolites and rishitin. The Δ*δ* values (in ppm) are defined as *δ* (metabolite)[ *δ* (rishitin), and the plus/minus values indicate low/high-field shifts compared to the shifts of rishitin, respectively. The shifts for the eliminated protons (H7 in 2 and H5 in 4) are temporarily set to 0 ppm.

### 3.3. Purification and structural analysis of rishitin metabolites produced after the incubation with B. cinerea

The supernatants of 500 µM rishitin solutions incubated with *B. cinerea* mycelia were extracted with ethyl acetate and subjected to reversed-phase HPLC separation, yielding six rishitin metabolites **1a, 1b, 2, 3, 4,** and **5** (Supplemental Fig. 1). The chemical structures of these metabolites were determined based on their molecular formulae obtained by high-resolution MS data and NMR analysis (Fig. 3, Table 1, Supplementary Figs. S2-S8, and Materials and methods).

The most polar metabolites **1a** and **1b** have the same molecular formulae (C_14_H_22_O_2_), but in contrast to rishitin (C_14_H_22_O_2_), possess two additional hydrogen and oxygen atoms. They appear to be stereoisomers because their ^1^H NMR data were quite similar (Table 1). The most striking difference of **1a**,**b** from rishitin in the NMR spectra is the lack of the 1,1-disubstituted alkene at C10-C11 and a newly emerging hydroxymethylene (-CH_2_OH) moiety, which resulted in the high-field shifts at H11 (Fig. 3). The hydroxymethylene moiety was unambiguously demonstrated by the change of multiplicity at H11 from two dd to two sharp d (AB quartet type) by D_2_O exchange experiment. The other structural parts were found to be the same as those for rishitin by 2-dimensional NMR experiments (DQF-COSY) (Supplementary Fig. S3 and 4). The small shift differences at H6-H8 (Fig. 3) could be due to the hydroxyl group at C10. These findings support the proposed structures of **1a**,**b**. The stereochemistry at C10 of each metabolite was not determined.

The metabolites **2**, **3**, and **5** are isomers possessing the molecular formula of C_14_H_22_O_3_, and exceed the mass of rishitin by one oxygen. Therefore, they were thought to be monooxygenated (hydroxy or epoxy) rishitins. A DQF-COSY experiment of **2** indicated the disconnection between the ethylidene H5-H6 and the methylene H8 due to the lack of methine H7 (Supplementary Fig. S5). This observation and the small low-field shift of the adjoining protons (H6, H8, and H11) indicate that **2** is 7-hydroxyrishitin (Fig. 3). The significant chemical shift difference in **3** from rishitin was the low-field shifts at H12 (Δδ = +2.4 ppm, Fig. 3). The DQF-COSY data indicated that other structural parts were identical to those of rishitin (Supplementary Fig. S6). These findings and small low-field shift at H11 indicate that **3** is 12-hydroxyrishitin (Fig. 3). The most characteristic feature of **5** in NMR is the appearance of a new olefinic proton (δ 5.66, -CH=), which was not observed for rishitin. A DQF-COSY experiment indicated that this olefinic proton connected to the H6-H7-H8 substructure (Supplementary Fig. S8). Since the two methylene protons H5 of rishitin disappeared in **5**, the new olefinic proton was assigned to H5. This finding suggested that the olefin C4a=C8a in rishitin was rearranged to C4a=C5. On the other hand, position C8a was possibly hydroxylated in view of the molecular formula of **5**. Therefore, **5** is not a simple hydroxy product but rather the 8a-hydroxylation accompanied by an olefin shift as shown in Fig. 3. This was supported by the significant low-field shifts at H5, small low-field shifts at H6, and small high-field shifts at H1 and H8 (Fig. 3). The stereochemistry of the introduced hydroxy group in **2** and **5** was not determined. Unlike the simple hydroxyrishitins, compound **4** was unique among the metabolites because it showed UV absorption at 244 nm, suggesting the presence of a conjugated ketone system in the molecule. A DQF-COSY experiment indicated the presence of the H6-H7-H8 connectivity but a lack of the signals for H5, suggesting that **4** is 5-ketorishitin or the dehydro product of 5-hydroxyrishitin. This was supported by the molecular formula smaller than those for the hydroxyrishitins by H_2_ and the low-field shifts at H4 and H6 due to the anisotropic effect (Fig. 3, Supplementary Fig. S7).

### 3.4. Purified 6 rishitin metabolites showed impaired antimicrobial activity against P. infestans and A. solani

To investigate the toxicity of the six purified rishitin metabolites, rishitin-sensitive *P. infestans* and *A. solani* were incubated with these compounds, to evaluate the effect of rishitin oxidation on antimicrobial activity. Mycelial growth of these two pathogens was examined after treatment with 500 µM of rishitin or its 6 metabolites. While 4a,8a-dihydro-4a,5-didehydro-8a-hydroxyrishitin (5) slightly inhibited the hyphal growth of pathogens, the other 5 metabolites lost their antimicrobial activity against *P. infestans* and *A. solani* (Fig. 4). Results with similar trends were obtained with the antimicrobial activity test using *A. brassicicola* (Supplemental Fig. S9). These results indicate, that all six compounds are indeed products of rishitin detoxification by *B. cinerea*.

**Fig. 4.**
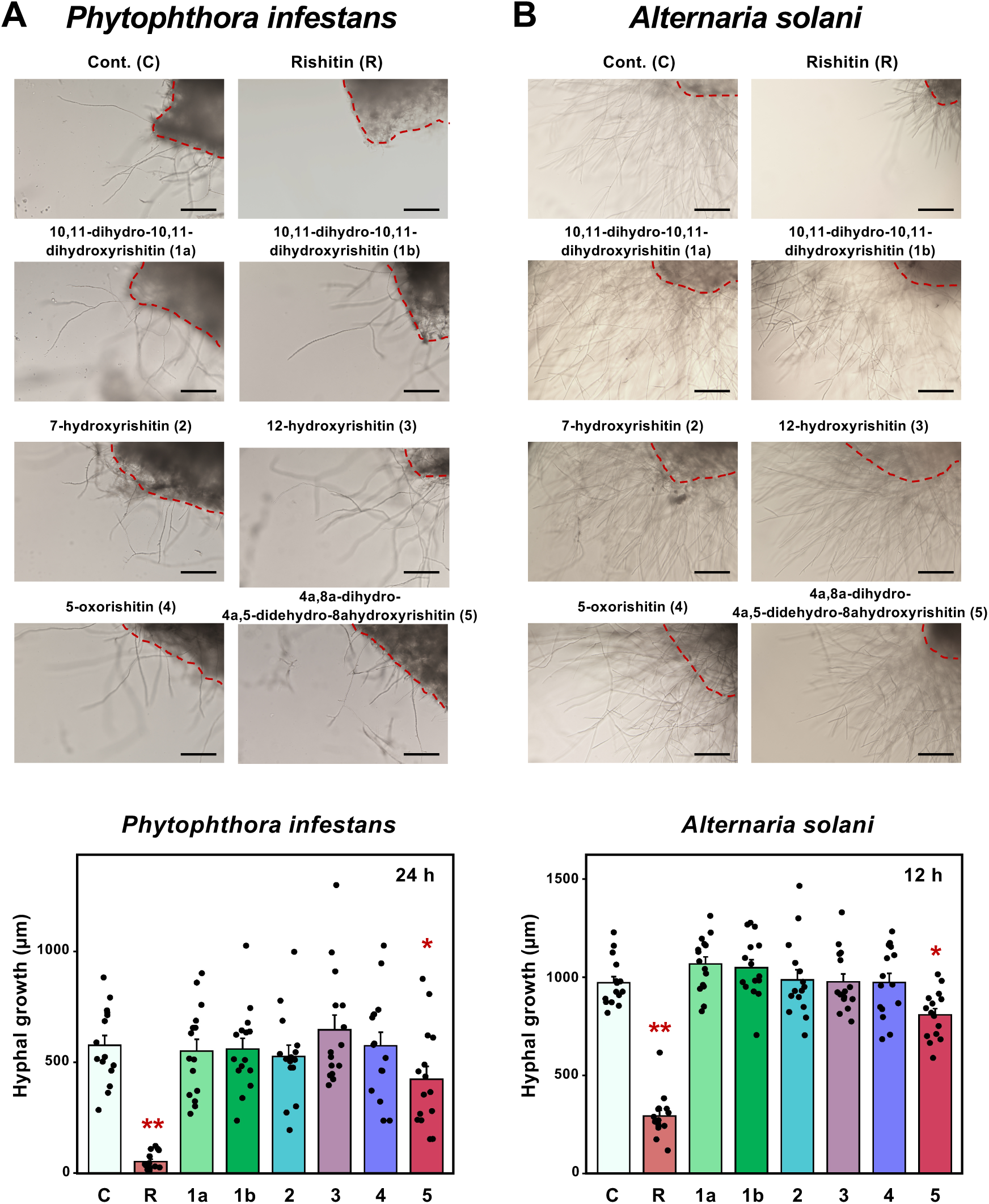
Anti-microbial activity of rishitin metabolites against phytopathogens. Mycelial blocks (approx. 1 mm^3^) of *Alternaria solani* **(A)** and *Phytophthora infestans* **(B)** were incubated in 50 µl of 0.1% DMSO (Cont. or C), 500 µM rishitin (R) or 500 µM rishitin metabolites. Outgrowth of hyphae from the mycelial block (outlined by dotted red lines) was measured after 12 or 24 h incubation. Bars = 200 µm. Data are means ± SE (n = 15). Data marked with asterisks are significantly different from the control as assessed by the two-tailed Student’s *t*-test: **P < 0.01, *P < 0.05.

### 3.5. Host-specialized phytopathogenic Botrytis species can partially metabolize rishitin

Fungi in the genus *Botrytis* include a number of phytopathogenic fungi. *B. cinerea*, and its cryptic species *B. pseudocinerea* and *B. prunorum*, are polyphagous pathogens, whereas most of *Botrytis* species have limited host range (Garfinkel et al., 2021). To compare the rishitin detoxification capacity of other *Botrytis* species, the metabolism of rishitin by four *Botrytis* species with narrow host range, namely *B. elliptica* (isolated from lily), *B. allii* (from onion), *B. squamosa* (from Chinese chive) and *B. tulipae* (from tulip), were investigated. As these *Botrytis* species are sensitive to rishitin compared with *B. cinerea* at 500 µM (Kuroyanagi et al., 2022), mycelial blocks of these species are incubated in 100 µM rishitin for 72 h.

As shown in Fig. 2, *B. cinerea* can completely metabolize rishitin into various metabolites within 72 h. Four host-specialized phytopathogenic *Botrytis* species also can metabolize rishitin, although remaining rishitin was detected for these species after 72 h incubation (Fig. 5 top), indicating that *B. cinerea* retains the highest rishitin metabolic capacity. Significant reduction in rishitin content was shown by *B. elliptica*, while *B. allii*, and *B. squamosa* and *B. tulipae* exhibited minimal reduction of rishitin. Comparison of the profiles of oxidized rishitin produced by the four species indicated that all tested species have similar rishitin metabolic pathways as *B. cinerea*, though the quantity ratios are variable (Fig. 5 middle). Similarly, these four species were also shown to be capable of metabolizing rishitin into two stereoisomers of 10,11-dihydro-10,11-dihydroxyrishitin (Fig. 5 bottom).

**Fig. 5.**
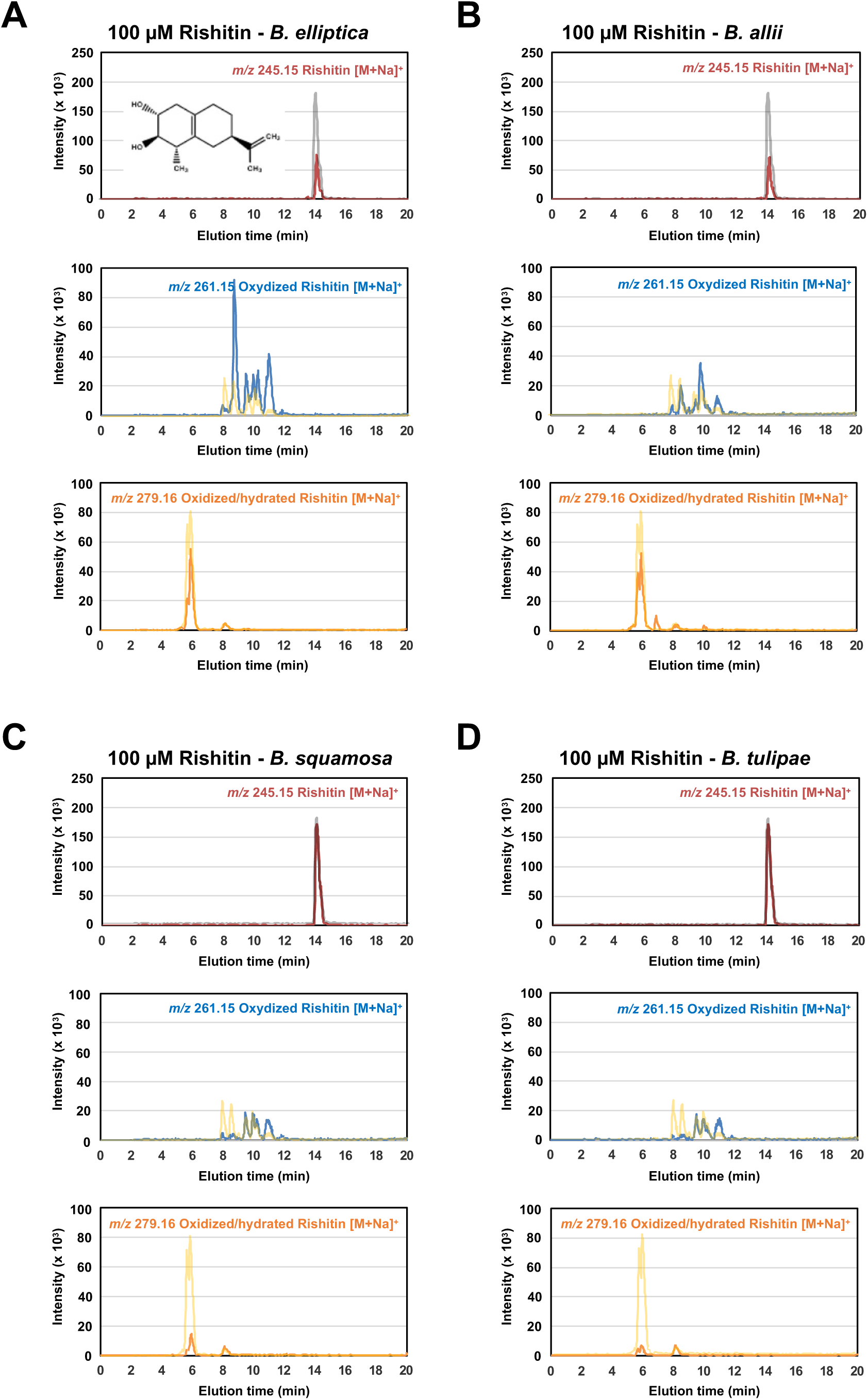
Metabolism of rishitin by *Botrytis* species. Mycelial block (approx. 1 mm^3^) of *Botrytis* species (**A,** *B. elliptica*; **B,** *B. allii*; **C,** *B. squamosa*; **D,** *B. tulipae*) was incubated in 100 µM rishitin for 72 h. Rishitin or its metabolites were detected by LC/MS. (top) Detection of rishitin (m/z 245.15) incubated with indicated *Botrytis* species. Profile of rishitin without incubation of mycelia is shown in grey lines. (middle and bottom) Detection of oxidized rishitins (middle, m/z 261.15) and oxidized/hydrated rishitins (bottom, m/z 279.16) after incubation with indicated *Botrytis* species. Profiles of metabolites after incubation with *B. cinerea* are shown in yellow lines.

## 4. Discussion

Induced production of phytoalexins, in response to pathogen attack, is a basal resistance mechanism commonly employed by plants against pathogens. Some polyxenous phytopathogens are known to detoxify phytoalexins enzymatically or export phytoalexins by transporters (Schoonbeek et al., 2001; Pedras and Ahiahonu, 2005; Sexton et al., 2009; Kuroyanagi et al., 2022; Bulasag et al, 2023). In contrast, biotrophic and hemi-biotrophic pathogens (typically *P. infestans*) generally use a vast array of effectors to suppress plant resistance mechanisms, including the production of phytoalexins (Zhou et al., 2011; Whisson et al., 2016). The toxicity of phytoalexins is often not specific to pathogens but is also harmful to plant cells (Shiraishi et al., 1975; Lyon, 1980; Stukkens et al., 2005). Thus, in plants, phytoalexins are effectively transported and exported to the site of pathogen attack, and accumulated phytoalexins are detoxified by healthy plant tissues (Shibata et al., 2016; Camagna et al., 2019; He et al., 2019).

### 4.1. Proposal of 5 pathways for the metabolism of rishitin by B. cinerea

The results of this study indicate that rishitin is converted into a variety of metabolites by *B. cinerea*. We propose five pathways for the detoxification of rishitin by *B. cinerea*, which we deem most likely to explain the observed compounds (Fig. 6). The rishitin metabolites determined in this study were all oxidized derivatives, possibly produced by monooxygenases (Fig. 3). 7-hydroxyrishitin (**2**) and 12-hydroxyrishitin (**3**) are both simple hydroxylation products, whereas the other metabolites could be produced by undergoing multi-step reactions including hydroxylation and epoxidation, etc. (Fig. 6). The isomeric metabolites 10,11-dihydro-10,11-dihydroxyrishitins (**1a** and **1b**) are possibly produced by the epoxidation at the C10-C11 double bond followed by the ring opening via a water molecule. It is unclear whether the second step (hydration) is enzymatic (Fig. 6). The unique ketone product 5-oxorishitin (**4**) could be produced by the hydroxylation at C5 followed by dehydrogenation of the resulting hydroxy group. Although the dehydrogenation of an allyl alcohol to the conjugated ketone is a common chemical reaction, a second enzyme such as dehydrogenase may be involved in the second reaction (Fig. 6). 4a,8a-dihydro-4a,5-didehydro-8a-hydroxyrishitin (**5**) is possibly produced by the epoxidation at the C4a-C8a double bond followed by the ring opening accompanied by H5 proton shift. It is also unclear whether a second enzyme is involved in the isomerization steps.

**Fig. 6.**
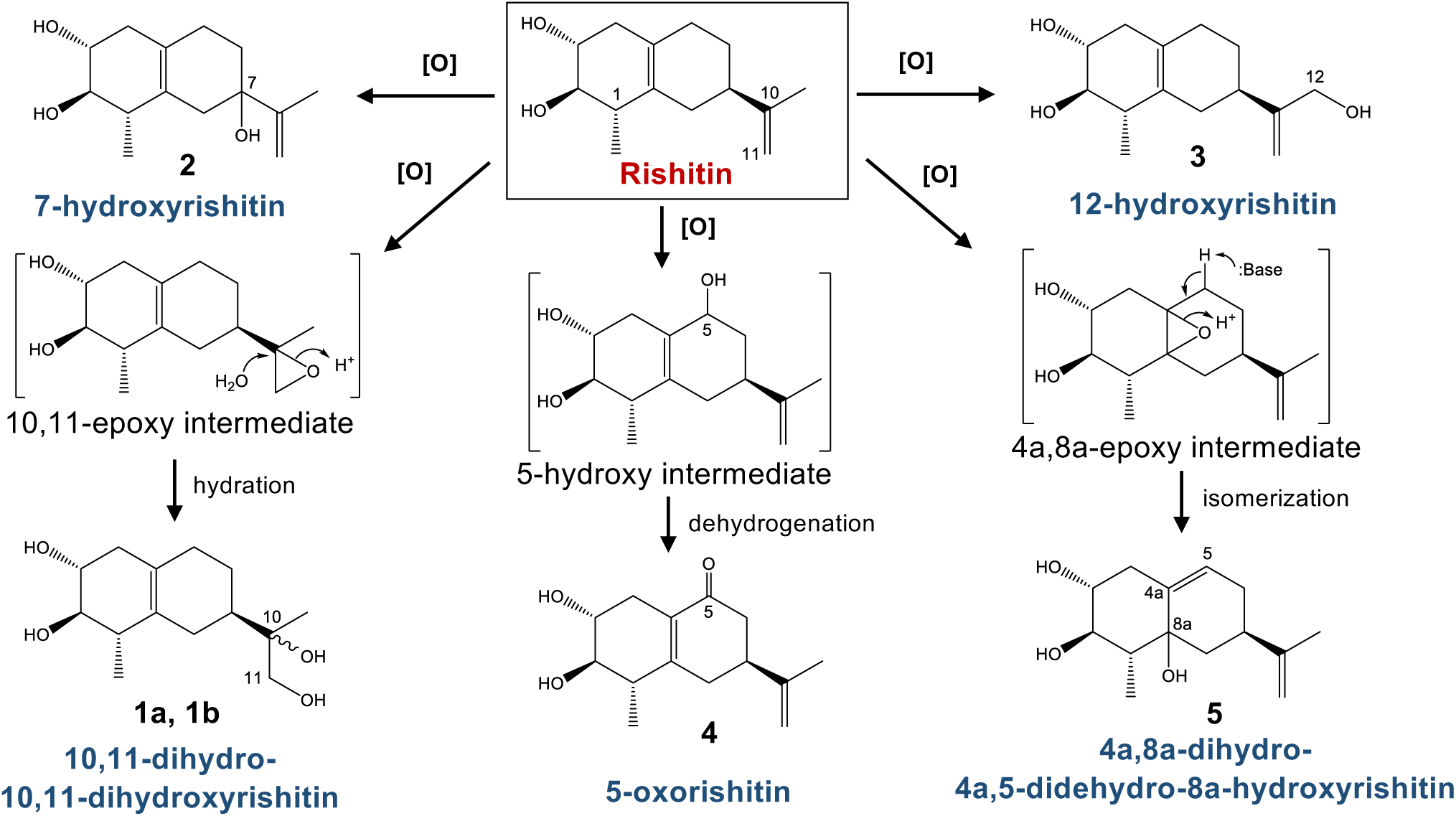
Predicted pathways for the metabolism of rishitin by *Botrytis cinerea.* **1a, 1b,** Rishitin is oxidized to form a tentative 10,11-epoxy intermediate that then undergoes hydration to form a 10,11-diols that are stereoisomeric at C10; **2,** Hydroxylation of rishitin at C7; **3,** Hydroxylation of the methyl group at C12; **4,** Hydroxylation at C5 leads to a tentative 5-hydroxy intermediate that subsequently undergoes dehydrogenation to form a carbonyl group at C5; **5,** Oxidation at the C4a-C8a alkene of rishitin to form a tentative epoxide intermediate, which then undergoes isomerization that leaves an allylic alcohol system at C5-C4a-C8a.

### 4.2. Detoxification of rishitin by plant pathogens and plants

There have been some previous reports on the detoxification of rishitin. *Gibberella pulicaris* (anamorph *Fusarium sambucinum*), the dry rot pathogen of potato tubers, oxidizes rishitin to 12-hydroxyrishitin and 10,11-epoxyrishitin (described as 13-hydroxyrishitin and 11,12-epoxyrishitin in Gardner et al. 1994). Rishitin is also metabolized by healthy potato tuber to less toxic rishitin M1 (12-hydroxyrishitin) and rishitin M2 (Ishiguri et al., 1978). A cytochrome P450 gene *StSPH* (sesquiterpenoid phytoalexins hydroxylase, a cytochrome P450), which metabolizes rishitin to rishitin M1, has been isolated from potato (Camagna et al., 2019). Homologous genes to *SPH* are exclusively found and widespread in the Solanaceae family, which produce sesquiterpenoid phytoalexins (Camagna et al., 2019). StSPH catalyzes the same detoxification reaction that occurs in *B. cinerea* and *G. pulicaris* (Fig. 6, Ishiguri et al., 1978; Gardner et al. 1994), but genes encoding enzymes homologous to plant SPH homologs were not found in the available fungal genome sequences including *B. cinerea* and *Fusarium* species. This result indicates that the host plants and phytopathogens independently evolved mechanisms to inhibit the antimicrobial activity of rishitin via detoxification to 12-hydroxyrishitin.

Previously, RNAseq analysis of upregulated genes in capsidiol-treated *B. cinerea* has allowed us to isolate and characterize capsidiol detoxifying enzyme genes *Bccpdh* (Bcin08g00930) and Bcin16g01490, by heterologous expression of candidate genes in grass endophytic fungi *Epichloë festucae* (Kuroyanagi et al., 2022). Further studies on the rishitin-responsive *B. cinerea* genes should allow us to isolate the genes involved in these detoxification reactions, to elucidate the evolutionary origin of rishitin detoxification in *B. cinerea*, and further help us understand why *B. cinerea* evolved five separate pathways for the detoxification of a single phytoalexin.

### 4.3. Metabolism of capsidiol and rishitin by Botrytis species

Capsidiol is the major sesquiterpenoid phytoalexin produced by *Nicotiana* and *Capsicum* species, which is metabolized to less toxic capsenone via dehydration by *B. cinerea* (Ward and Stoessl, 1972, Kuroyanagi et al., 2022). Metabolism of capsidiol by *B. elliptica*, *B. allii*, *B. squamosa,* and *B. tulipae* was not detected and the gene for capsidiol dehydrogenase *Bccpdh* was exclusively found in *B. cinerea*, but not in the genome of the other *Botrytis* species. Comparison of the genome sequences of *B. cinerea* and its closely related species suggested that *B. cinerea* may have acquired *Bccpdh* by means of horizontal gene transfer (Kuroyanagi et al., 2022). In contrast, the results of this study indicated that in *Botrytis* species, rishitin metabolism is, albeit to varying degrees, widespread, despite the fact that the four host-specific *Botrytis* species we analyzed do not infect rishitin producing plants. It follows therefore that the ability to metabolize rishitin is likely to be an ancestral trait, which is conserved in *Botrytis* species.

In the previous study, the ABC transporter *BcatrB* was shown to be essential for rishitin resistance and infection of plants producing rishitin, suggesting that it may act as a rishitin efflux pump (Bulasag et al., 2023). BcatrB has also been involved in the export of structurally unrelated phytoalexins such as resveratrol, camalexin (Vermeulen et al., 2001; Stefanato et al., 2009), and phenylpyrrole fungicides (Schoonbeek et al., 2001), thus BcatrB is considered as a multidrug resistance transporter with low substrate specificity. Homologs of *BcatrB* are also conserved in the *Botrytis* genus (Bulasag et al., 2023). It has been found that fungicide-resistant field populations of *B. cinerea* show significantly increased basal expression of *BcatrB* (Kretschmer et al., 2009), indicating that evolving a more efficient efflux of newly encountered harmful substances appears to be an effective first measure to tolerate toxins, and seems to evolve faster than a specific detoxification pathway. It might therefore be the case that *B. cinerea* acquired its broad host-range by means of an efficient phytoalexin efflux, allowing it to establish a beachhead on a new host, before further adapting via more sophisticated detoxification mechanisms. Discovery of the genes involved in the rishitin detoxification may allow further insights into how efflux and detoxification regulation are intertwined in *B. cinerea*, and shed light on the evolutionary history that sets it apart from the various host-specific *Botrytis* species.

## Author contributions

DT designed the research. ASB, AA, AM, SP, TK, MO, and DT conducted the experiments. ASB, MC, MO, and DT analyzed data. AT, IS, SC, MO, and DT supervised the experiments. ASB, AA, MC, AT, IS, SC, MO, and DT contributed to the discussion and interpretation of the results. ASB, MO, and DT wrote the manuscript. ASB, MC, MO, and DT edited the manuscript.

## Data availability

Raw data is available on request from the corresponding author, D.T.

## Acknowledgments

We are grateful to Drs. Kenji Asano and Kotaro Akai (National Agricultural Research Center for Hokkaido Region, Japan) and Mr. Yasuki Tahara (Nagoya University, Japan) for providing tubers of potato cultivars, and Ms. Kayo Shirai (Hokkaido Central Agricultural Experiment Station, Japan) and Dr. Seishi Akino (Hokkaido University, Japan) for providing *P. infestans* isolate 08YD1. We would like to acknowledge the Japanese Ministry of Education, Culture, Sports and Technology (MEXT) and the University of the Philippines Los Baños for allowing Abriel Salaria Bulasag to pursue graduate studies in Japan on scholarship. This work was supported by a Grant-in-Aid for Scientific Research (B) (20H02985 and 23H02212) and Grant-in-Aid for Challenging Exploratory Research (22K19176) to DT from the Japan Society for the Promotion of Science.

## Supplementary Figures

**Supplementary Fig. S1.** HPLC separation of rishitin metabolites.

**Supplementary Fig. S2.** LC/MS data of an extract of *B. cinerea* cultured in the presence of rishitin.

**Supplementary Fig. S3.** NMR of metabolite **1a** (in CDCl_3_ at 400 MHz).

**Supplementary Fig. S4.** NMR of metabolite **1b** (in CDCl_3_ at 400 MHz).

**Supplementary Fig. S5.** NMR of metabolite **2** (in CDCl_3_ at 400 MHz).

**Supplementary Fig. S6.** NMR of metabolite **3** (in CDCl_3_ at 400 MHz).

**Supplementary Fig. S7.** NMR of metabolite **4** (in CDCl_3_ at 400 MHz).

**Supplementary Fig. S8.** NMR of metabolite **5** (in CDCl_3_ at 400 MHz).

**Supplementary Fig. S9.** Anti-microbial activity of rishitin metabolites against phytopathogenic fungi *Alternaria brassicicola*.

## References

1. Ahuja, I., Kissen, R., Bones, A.M., 2012. Phytoalexins in defense against pathogens. Trends Plant Sci. 17, 73–90.

2. Allan, J., Regmi, R., Denton-Giles, M., Kamphuis, L.G., Derbyshire, M.C., 2019. The host generalist phytopathogenic fungus *Sclerotinia sclerotiorum* differentially expresses multiple metabolic enzymes on two different plant hosts. Sci. Rep. 9, 19966.

3. Bulasag, A.S., Camagna, M., Kuroyanagi, T., Ashida, A., Ito, K., Tanaka, A., Sato, I., Chiba, S., Ojika, M., Takemoto, D., 2023. *Botrytis cinerea* tolerates phytoalexins produced by Solanaceae and Fabaceae plants through an efflux transporter BcatrB and metabolizing enzymes. Front Plant Sci. 14, 1177060.

4. Camagna, M., Ojika, M., Takemoto D., Detoxification of the solanaceous phytoalexins rishitin, lubimin, oxylubimin and solavetivone via a cytochrome P450 oxygenase. Plant Signal. Behav. 15, 1707348.

5. Coleman, J.J., Wasmann, C.C., Usami, T., White, G.J., Temporini, E.D., McCluskey, K., VanEtten, H.D., 2011. Characterization of the gene encoding pisatin demethylase (FoPDA1) in *Fusarium oxysporum*. Mol. Plant Microbe Interact. 24, 1482–1491.

6. De Wit, P.J.G.M., Flach, W., 1979. Differential accumulation of phytoalexins in tomato leaves but not in fruits after inoculation with virulent and avirulent races of *Cladosporium fulvum*. Physiol. Plant Pathol. 15, 257–267.

7. Gardner, H.W., Desjardins, A.E., McCormick, S.P., Weisleder, D., 1994. Detoxification of the potato phytoalexin rishitin by *Gibberella pulicaris*. Phytochemistry 37, 1001–1005.

8. Garfinkel, A.R., 2021. The history of *Botrytis* taxonomy, the rise of phylogenetics, and implications for species recognition. Phytopathology 111, 437–454.

9. Hammerschmidt, R., 1999. Phytoalexins: What have we learned after 60 years? Annu. Rev. Phytopathol. 37, 285–306.

10. He, Y., Xu, J., Wang, X., He, X., Wang, Y., Zhou, J., Zhang, S., Meng, X., 2019. The Arabidopsis pleiotropic drug resistance transporters PEN3 and PDR12 mediate camalexin secretion for resistance to *Botrytis cinerea*. Plant Cell 31, 2206–2222.

11. Imano, S., Fushimi, M., Camagna, M., Tsuyama-Koike, A., Mori, H., Ashida, A., Tanaka, A., Sato, I., Chiba, S., Kawakita, K., Ojika, M., Takemoto, D., 2022. AP2/ERF transcription factor NbERF-IX-33 is involved in the regulation of phytoalexin production for the resistance of *Nicotiana benthamiana* to *Phytophthora infestans*. Front. Plant Sci. 12, 821574.

12. Ishiguri, Y., Tomiyama, K., Murai, A., Katsui, N., Masamune, T., 1978. Toxicity of rishitin, rishitin-M-1, and rishitin-M-2 to *Phytophthora infestans* and potato tissue. Ann. Phytopath. Soc. Japan 44, 52–56.

13. Katsui, N., Mural, A., Takasugi, M., Imaizumi, K., Masamune, T., Tomiyama, K., 1968. The structure of rishitin, a new antifungal compound from diseased potato tubers. Chem. Commun. 1, 43–44.

14. Kretschmer, M., Leroch, M., Mosbach, A., Walker, A.S., Fillinger, S., Mernke, D., Schoonbeek, H.J., Pradier, J.M., Leroux, P., De Waard, M.A., Hahn, M., 2009. Fungicide-driven evolution and molecular basis of multidrug resistance in field populations of the grey mould fungus *Botrytis cinerea*. PLoS Pathog. 5, e1000696.

15. Kuroyanagi, T., Bulasag, A.S., Fukushima, K., Ashida, A., Suzuki, T., Tanaka, A., Camagna, M., Sato, I., Chiba, S., Ojika, M., Takemoto, D., 2022. *Botrytis cinerea* identifies host plants via the recognition of antifungal capsidiol to induce expression of a specific detoxification gene. PNAS Nexus 1, pgac274.

16. Kusch, S., Larrouy, J., Ibrahim, H.M.M., Mounichetty, S., Gasset, N., Navaud, O., Mbengue, M., Zanchetta, C., Lopez-Roques, C., Donnadieu, C., Godiard, L., Raffaele, S., 2022. Transcriptional response to host chemical cues underpins the expansion of host range in a fungal plant pathogen lineage. ISME J. 16, 138–148.

17. Lyon, G.D., 1980. Evidence that the toxic effect of rishitin may be due to membrane damage. J. Exp. Bot. 37, 957–966.

18. Maude, R.B., Humpherson-Jones, F.M., 1980. Studies on the seed-borne phases of dark leaf spot (*Alternaria brassicicola*) and grey leaf spot (*Alternaria brassicae*) of brassicas. Ann. Appl. Biol. 95, 311–319.

19. Murai, A., Nishizakura, K., Katsui, N., Masamune, T., 1975. The synthesis of rishitin. Tetrahedron Lett. 16, 4399–4402.

20. Newman, T.E., Derbyshire, M.C., 2020. The evolutionary and molecular features of broad host-range necrotrophy in plant pathogenic fungi. Front. Plant Sci. 11, 591733.

21. Pedras, M.S.C., Abdoli, A., 2017. Pathogen inactivation of cruciferous phytoalexins: detoxification reactions, enzymes and inhibitors. RSC Adv. 7, 23633–23646.

22. Pedras, M.S., Ahiahonu, P.W., 2005. Metabolism and detoxification of phytoalexins and analogs by phytopathogenic fungi. Phytochemistry 66, 391– 411.

23. Prasad, L., Katoch, S., Shahid, S., 2022. Microbial interaction mediated programmed cell death in plants. 3 Biotech. 12, 43.

24. Sanchez-Vallet, A., Ramos, B., Bednarek, P., López, G., Piślewska-Bednarek, M., Schulze-Lefert, P., Molina, A., 2010. Tryptophan-derived secondary metabolites in *Arabidopsis thaliana* confer non-host resistance to necrotrophic *Plectosphaerella cucumerina* fungi. Plant J. 63, 115–127.

25. Schneider, C.A., Rasband, W.S., Eliceiri, K.W., 2012. NIH Image to ImageJ: 25 years of image analysis. Nat. Methods 9, 671–675.

26. Schoonbeek, H., Del Sorbo, G., De Waard, M.A., 2001. The ABC transporter BcatrB affects the sensitivity of *Botrytis cinerea* to the phytoalexin resveratrol and the fungicide fenpiclonil. Mol. Plant Microbe Interact. 14, 562–571.

27. Sexton, A.C., Minic, Z., Cozijnsen, A.J., Pedras, M.S., Howlett, B.J., Cloning, purification and characterisation of brassinin glucosyltransferase, a phytoalexin-detoxifying enzyme from the plant pathogen *Sclerotinia sclerotiorum*. Fungal Genet. Biol. 46, 201–209.

28. Shibata, Y., Kawakita, K., Takemoto, D., 2010. Age-related resistance of *Nicotiana benthamiana* against hemibiotrophic pathogen *Phytophthora infestans* requires both ethylene- and salicylic acid-mediated signaling pathways. Mol. Plant Microbe. Interact. 23.1130–1142.

29. Shibata, Y., Ojika, M., Sugiyama, A., Yazaki, K., Jones, D.A., Kawakita, K., Takemoto, D., 2016. The full-size ABCG transporters Nb-ABCG1 and Nb-ABCG2 function in pre- and postinvasion defense against *Phytophthora infestans* in *Nicotiana benthamiana*. Plant Cell 28, 1163–1181.

30. Shiraishi, T., Oku, H., Isono, M., Ouchi, S., 1975. The injurious effect of pisatin on the plasma membrane of pea. Plant Cell Physiol. 16, 939–942.

31. Stefanato, F.L., Abou-Mansour, E., Buchala, A., Kretschmer, M., Mosbach, A., Hahn, M., Bochet, C.G., Métraux, J.P., Schoonbeek, H.J., 2009. The ABC transporter BcatrB from *Botrytis cinerea* exports camalexin and is a virulence factor on *Arabidopsis thaliana*. Plant J. 58, 499–510.

32. Stukkens, Y., Bultreys, A., Grec, S., Trombik, T., Vanham, D., Boutry, M., 2005. NpPDR1, a pleiotropic drug resistance-type ATP-binding cassette transporter from *Nicotiana plumbaginifolia*, plays a major role in plant pathogen defense. Plant Physiol. 139, 341–352.

33. van Wees, S.C., Chang, H.S., Zhu, T., Glazebrook, J. 2003. Characterization of the early response of Arabidopsis to *Alternaria brassicicola* infection using expression profiling. Plant Physiol. 132, 606–617.

34. Vermeulen, T., Schoonbeek, H., De Waard, M.A., 2001. The ABC transporter BcatrB from *Botrytis cinerea* is a determinant of the activity of the phenylpyrrole fungicide fludioxonil. Pest Manag. Sci. 57, 393–402.

35. Ward, E.W.B., Stoessl, A., 1972. Postinfectional inhibitors from plants. III. Detoxification of capsidiol, an antifungal compound from peppers. Phytopathology 62, 1186.

36. Wasmann, C.C., VanEtten, H.D., 1996. Transformation-mediated chromosome loss and disruption of a gene for pisatin demethylase decrease the virulence of *Nectria haematococca* on pea. Mol. Plant Microbe. Interact. 9, 793–803.

37. Westrick, N.M., Smith, D.L., Kabbage, M., 2021. Disarming the host: detoxification of plant defense compounds during fungal necrotrophy. Front. Plant Sci. 12, 651716.

38. Whisson, S.C., Boevink, P.C., Wang, S., Birch, P.R., 2016. The cell biology of late blight disease. Curr. Opin. Microbiol. 34, 127–135.

39. Zhou, H., Lin, J., Johnson, A., Morgan, R.L., Zhong, W., Ma, W., 2011. *Pseudomonas syringae* type III effector HopZ1 targets a host enzyme to suppress isoflavone biosynthesis and promote infection in soybean. Cell Host Microbe. 9, 177–186.

