## Supplementary Figures for "*Botrytis cinerea* detoxifies the sesquiterpenoid phytoalexin rishitin through multiple metabolizing pathways"

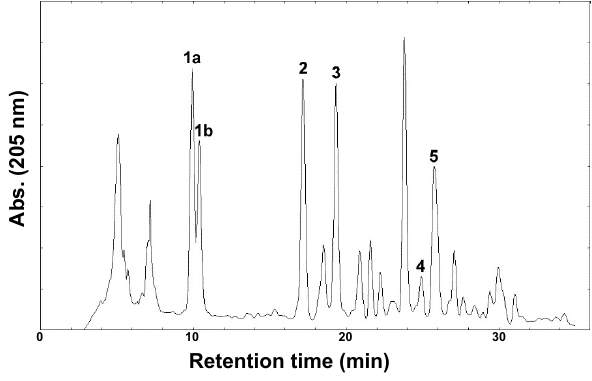

**Supplementary Fig. S1.** HPLC separation of rishitin metabolites **1a**, **1b**, **2**, **3**, **4**, and **5.** The peak at 25.8 min was further purified under different conditions to obtain pure **5**.

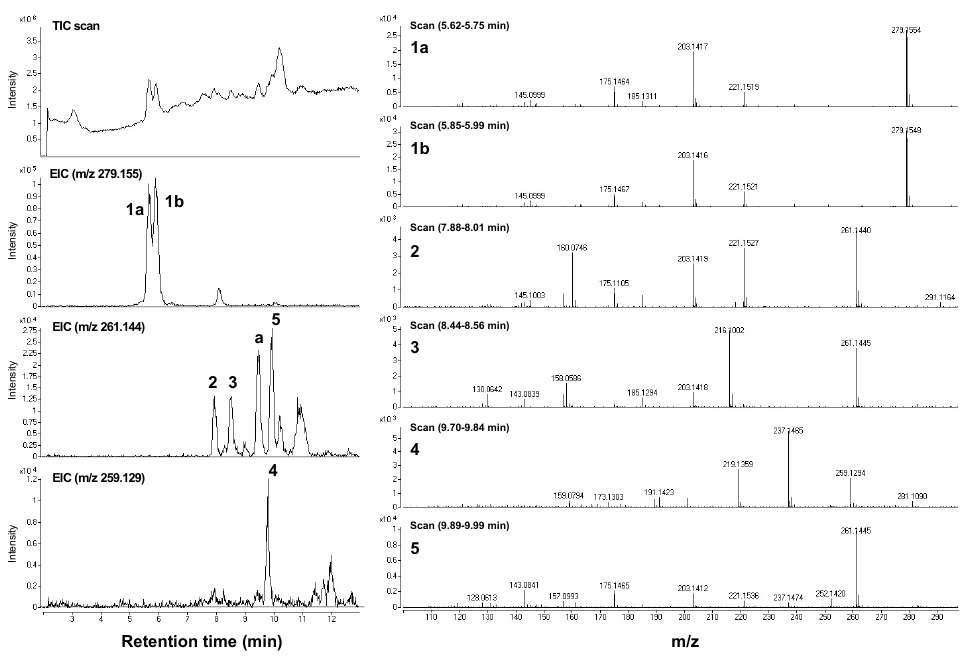

**Supplementary Fig. S2.** LC/MS data of an extract of *B. cinerea* cultured in the presence of rishitin for 72 h. Left: LC/MS data of an extract of *B. cinerea* cultured in the presence of rishitin. Left: TIC (top) and extracted ion chromatographs. The peak “a” was not characterized due to a trace yield in contrast with the peak intensity. Right: MS of the identified metabolites.

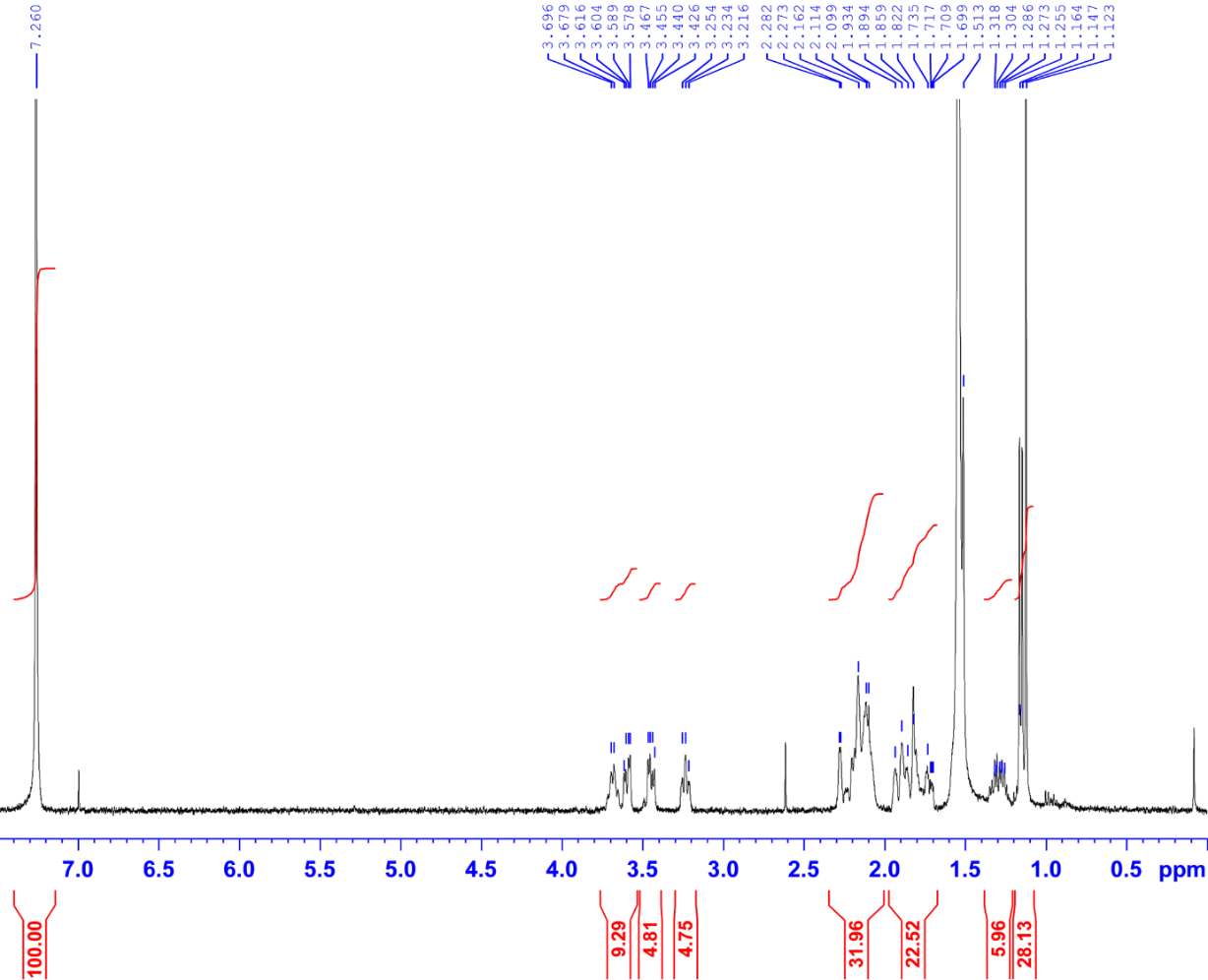

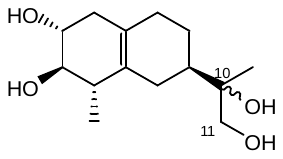

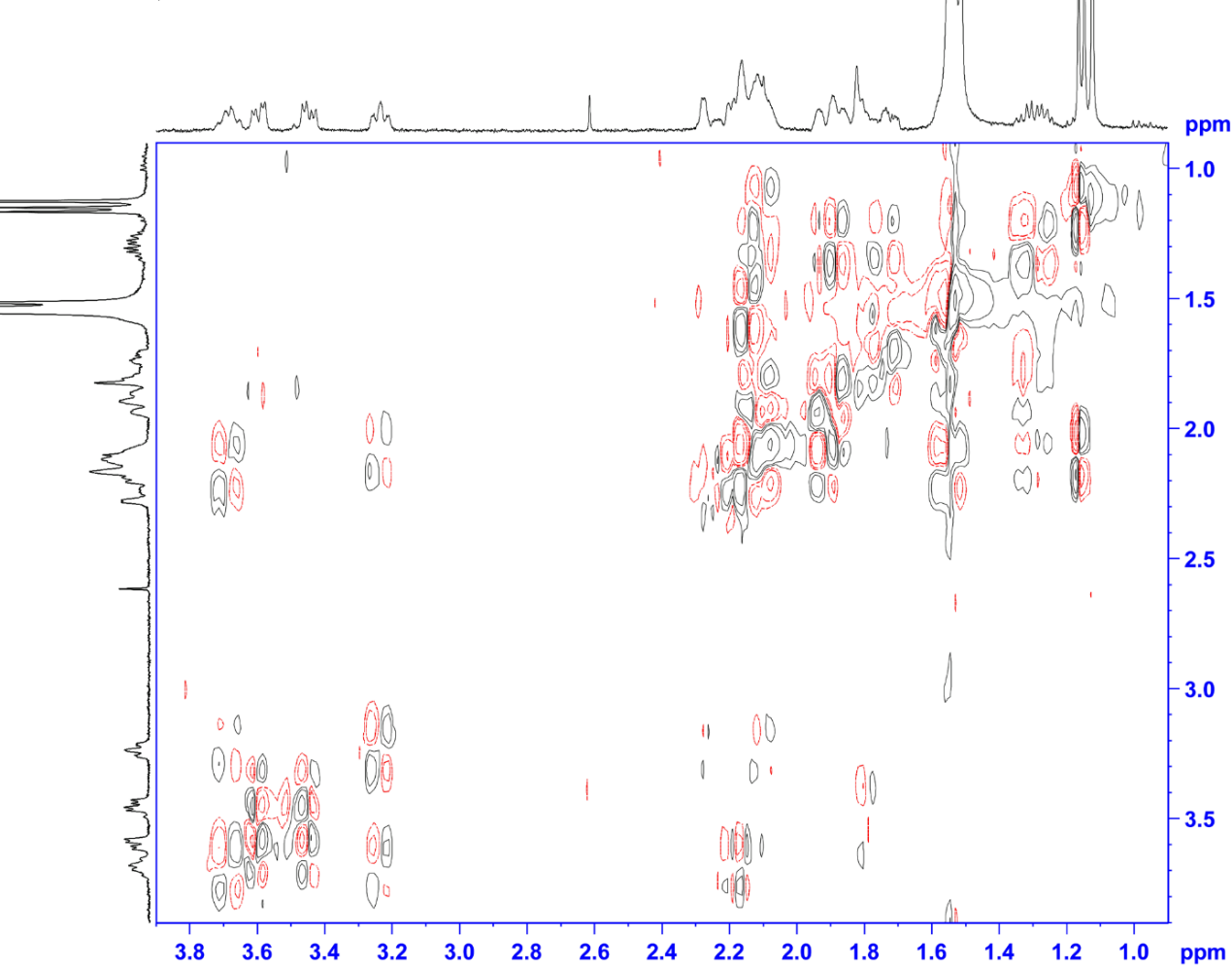

**Supplementary Fig. S3.** NMR of metabolite **1a** (in CDCl_3_ at 400 MHz). (from top) ^1^H NMR, DQF-COSY.

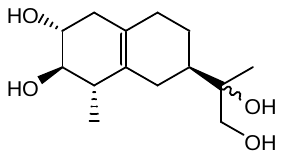

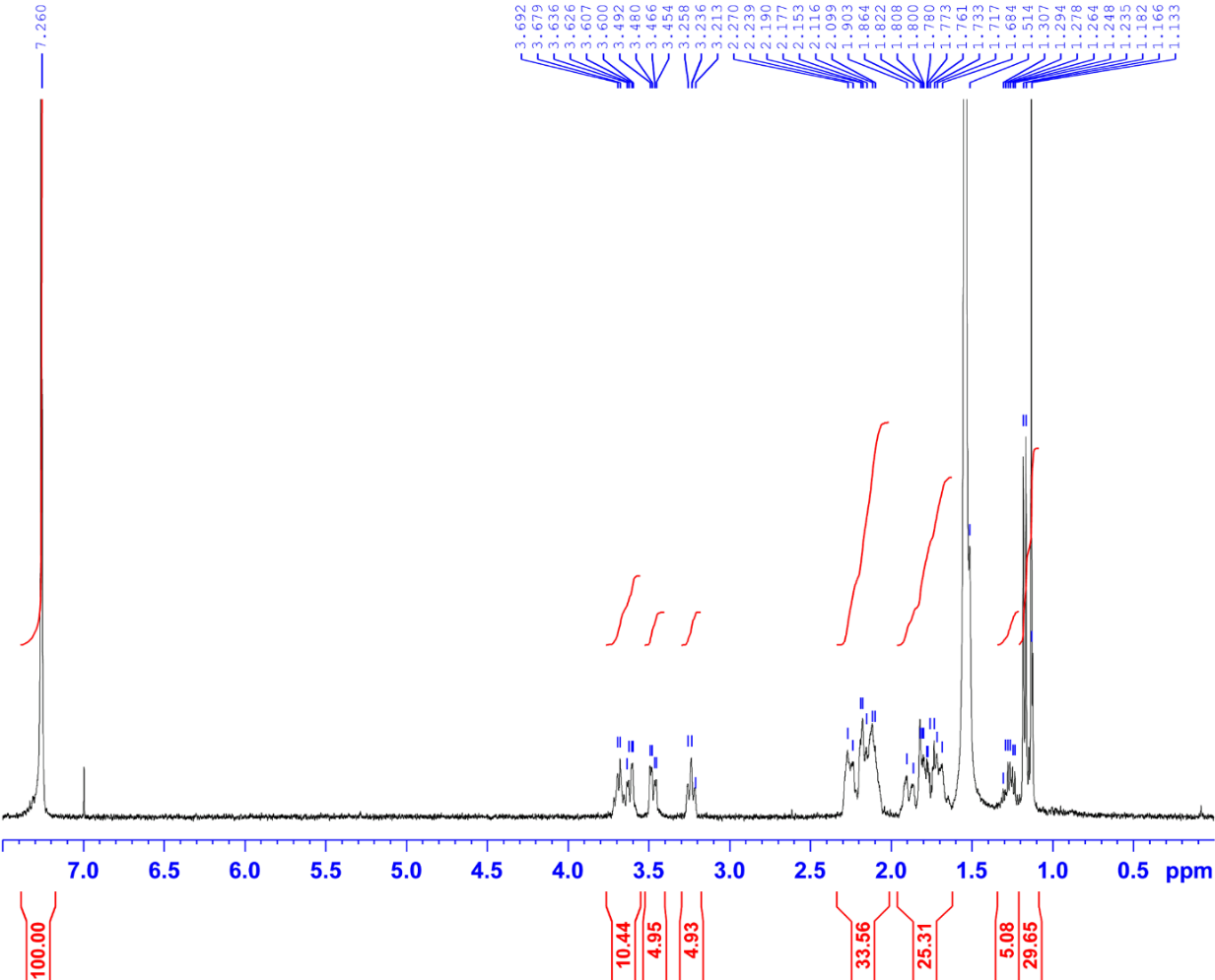

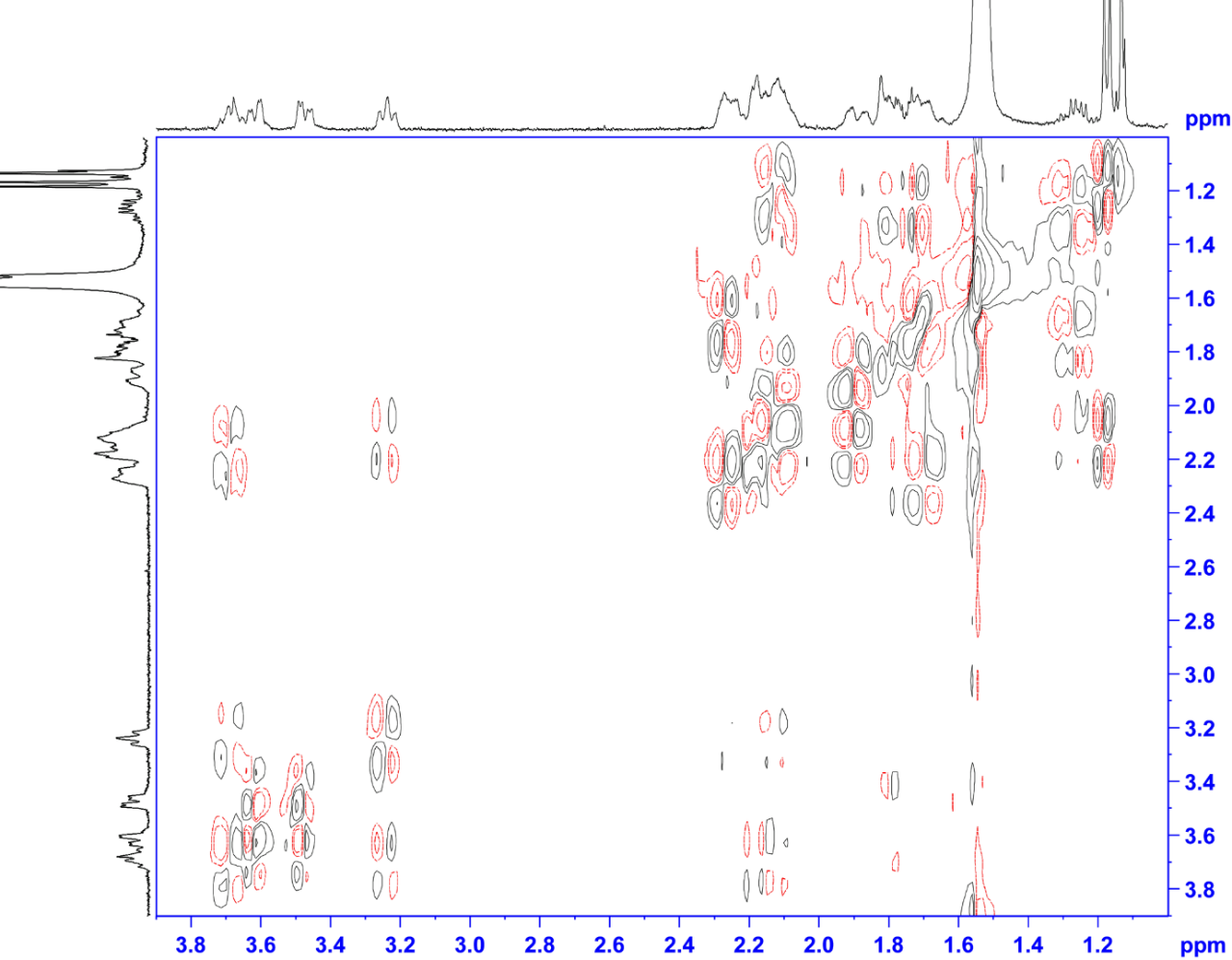

**Supplementary Fig. S4.** NMR of metabolite **1b** (in CDCl_3_ at 400 MHz). (from top) ^1^H NMR, DQF-COSY.

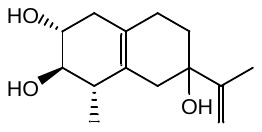

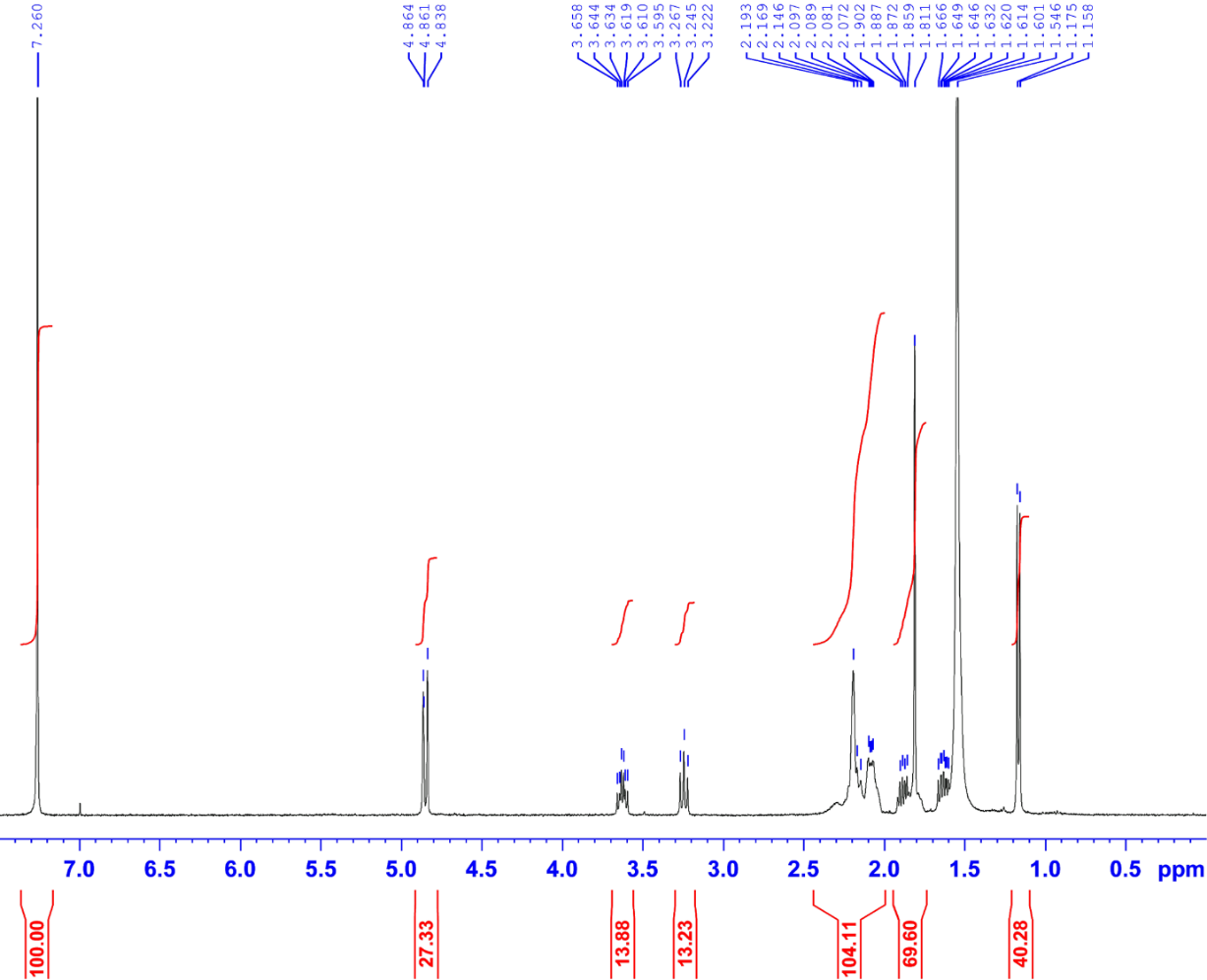

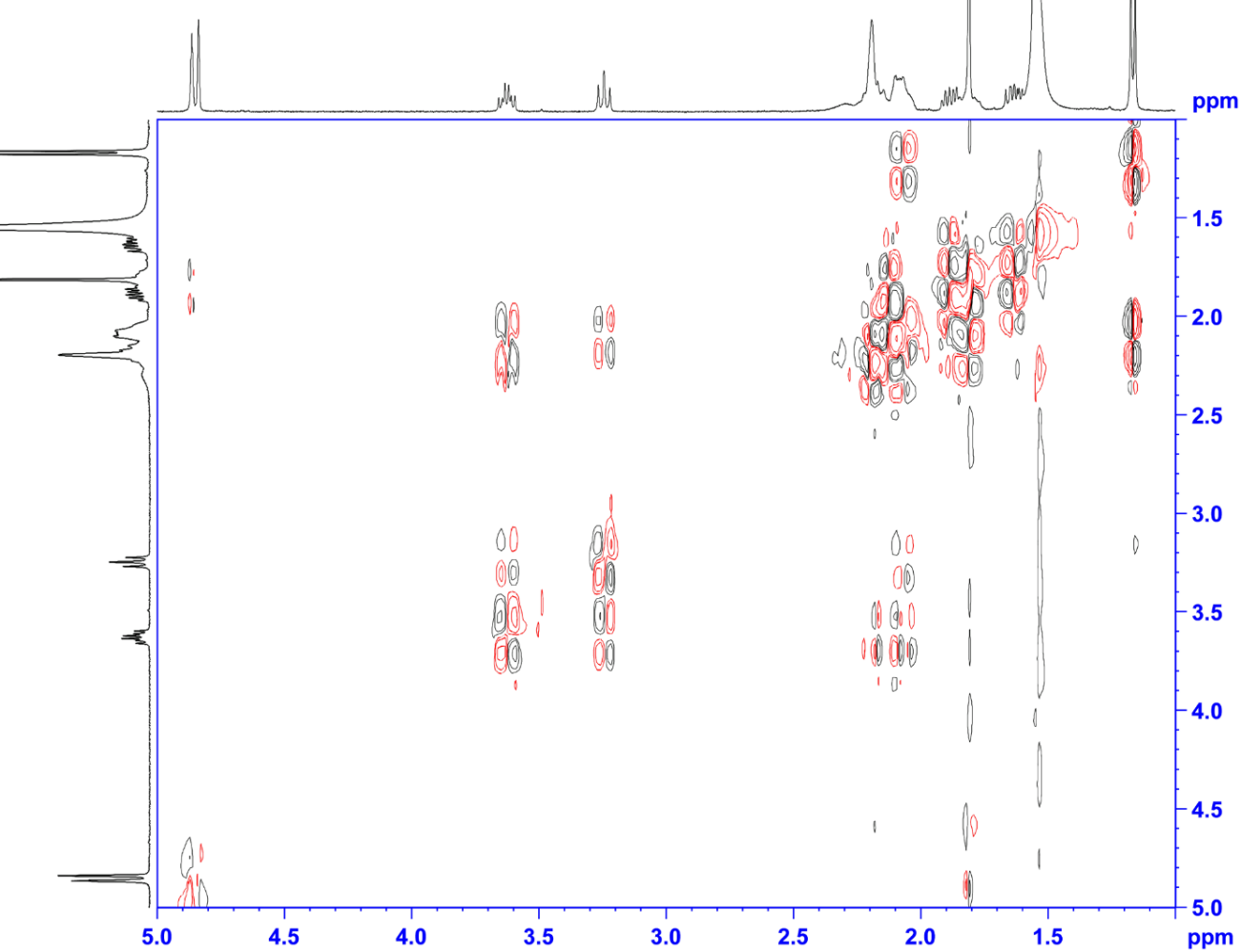

**Supplementary Fig. S5.** NMR of metabolite **2** (in CDCl_3_ at 400 MHz). (from top) ^1^H NMR, DQF-COSY.

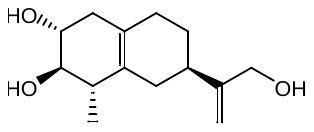

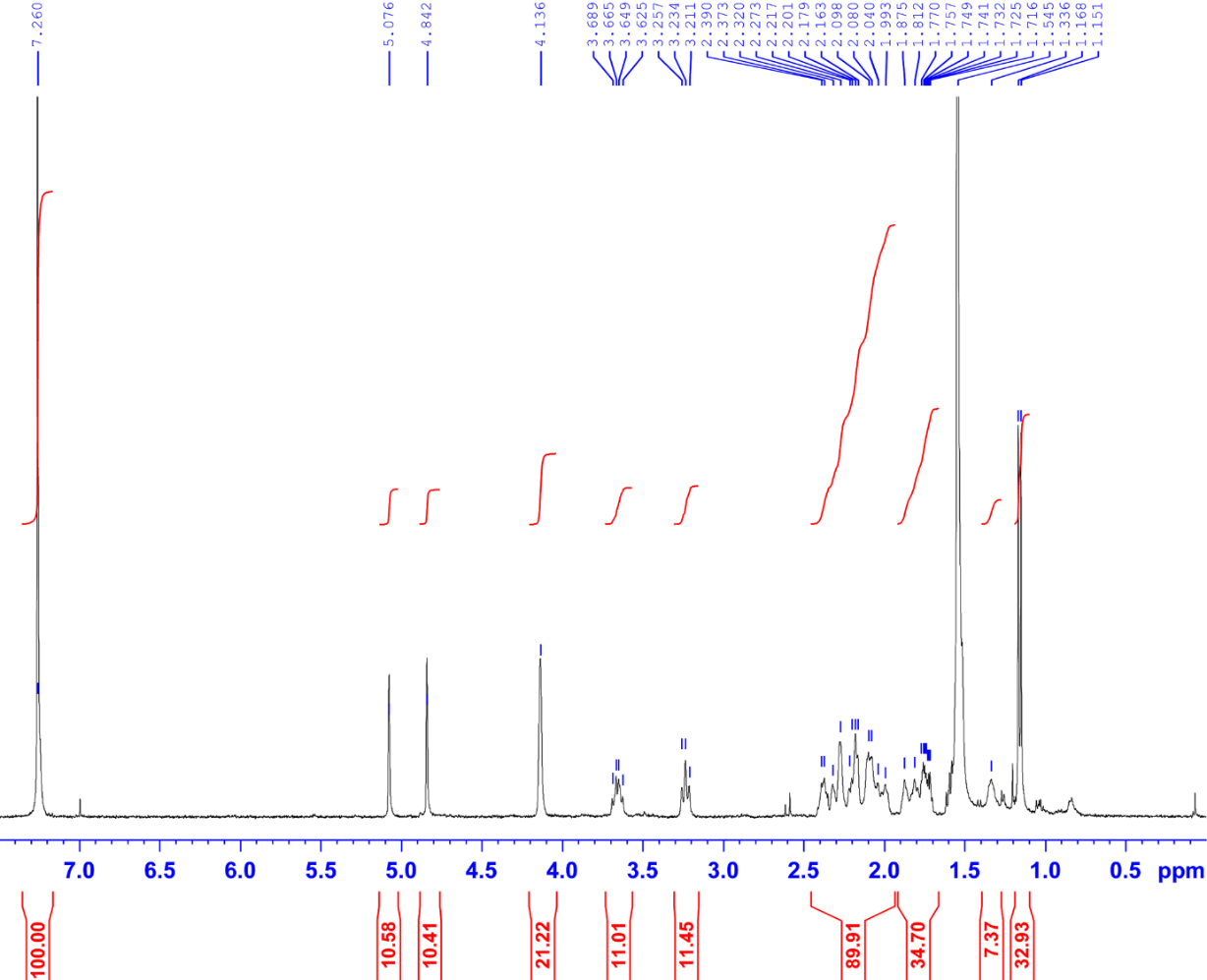

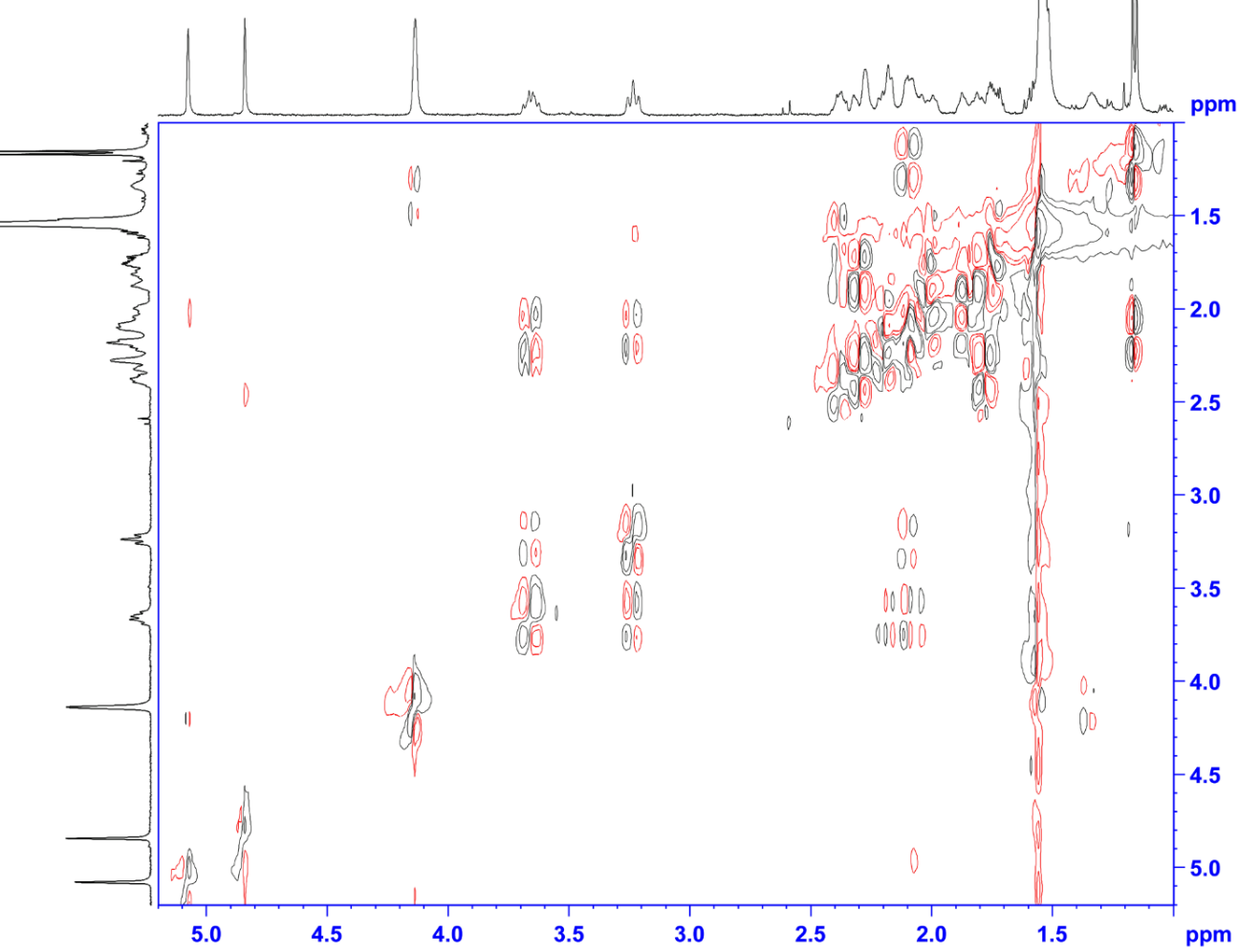

**Supplementary Fig. S6.** NMR of metabolite **3** (in CDCl_3_ at 400 MHz). (from top) ^1^H NMR, DQF-COSY.

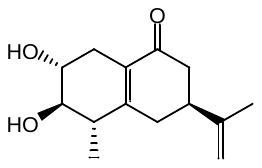

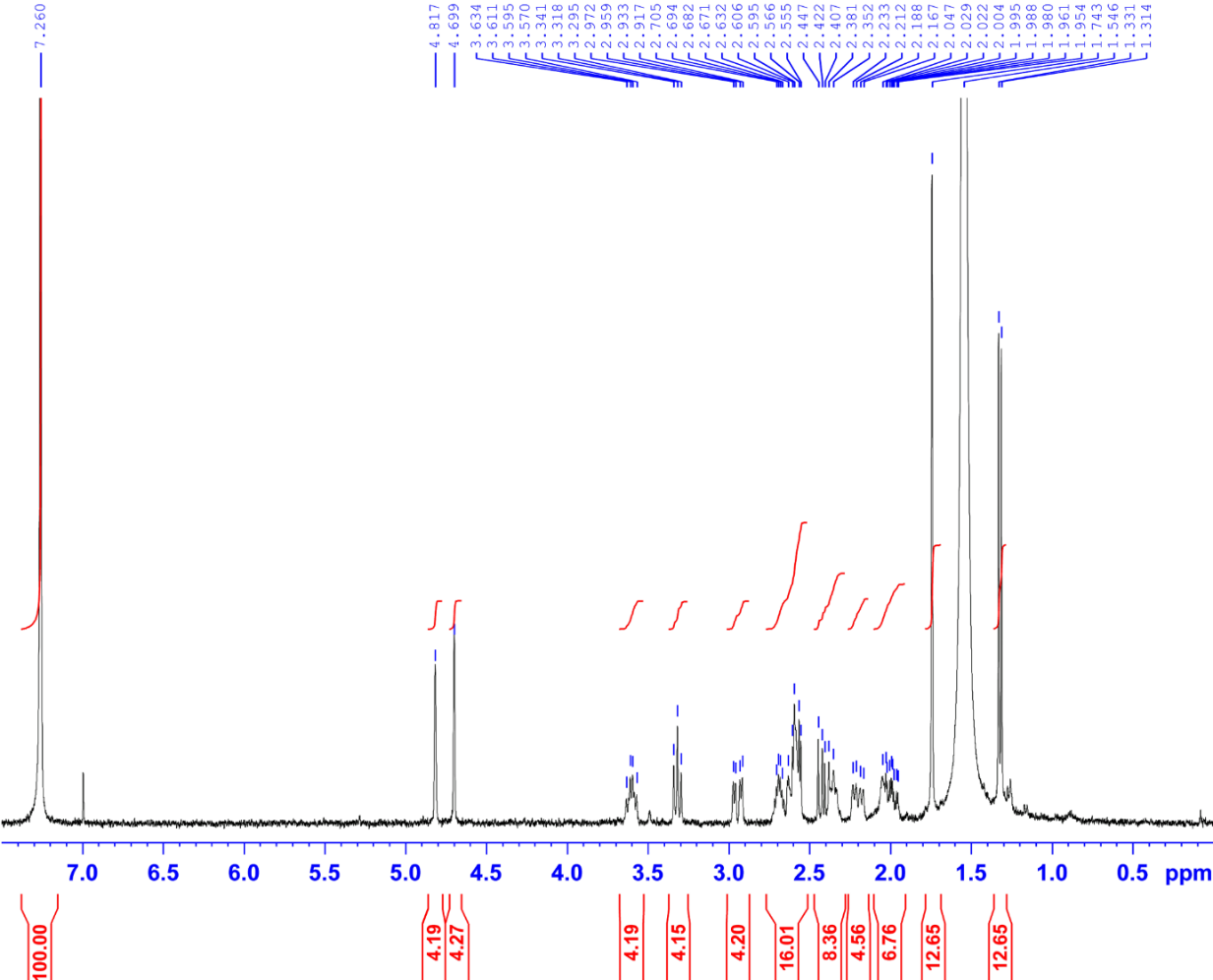

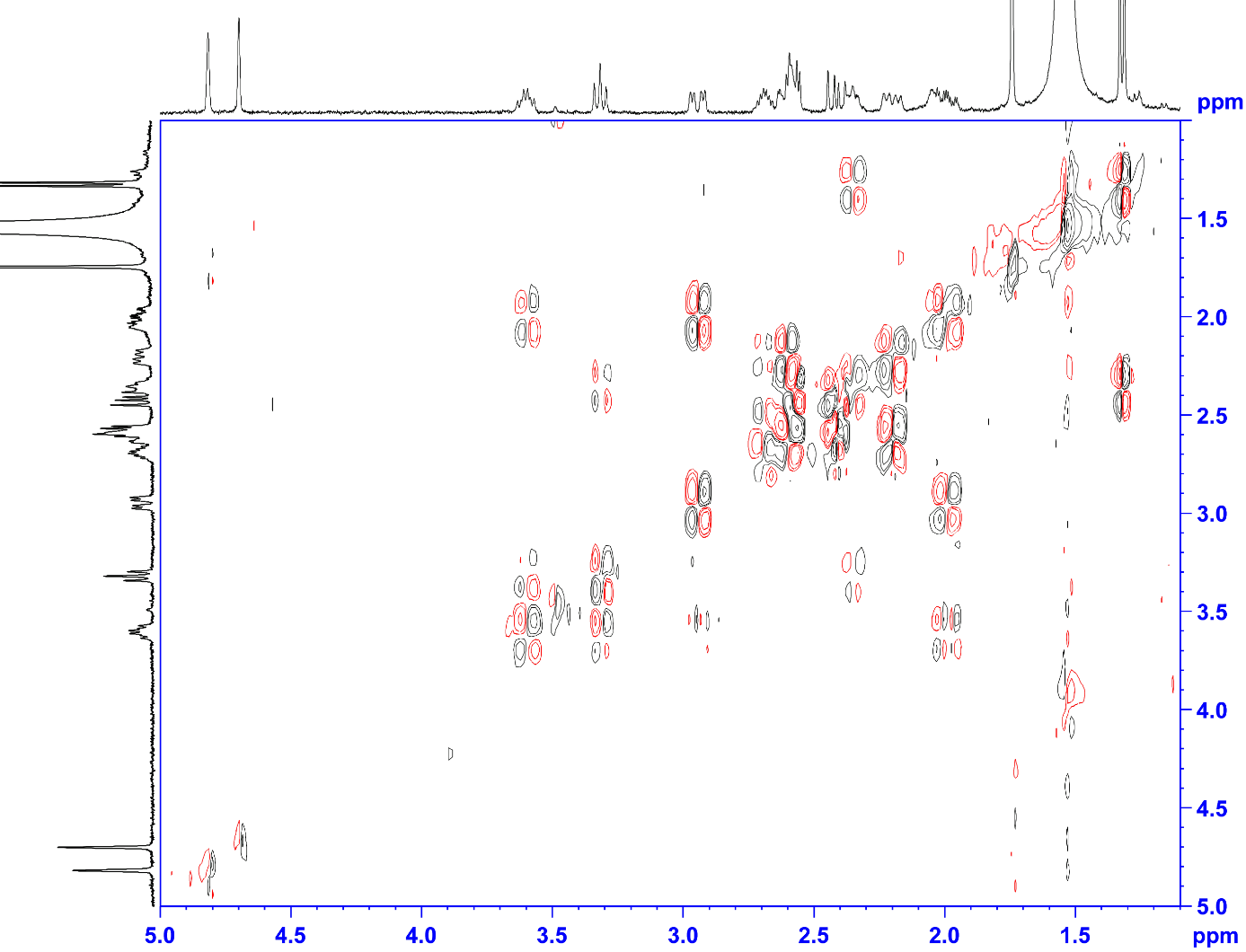

**Supplementary Fig. S7.** NMR of metabolite **4** (in CDCl_3_ at 400 MHz). (from top) ^1^H NMR, DQF-COSY.

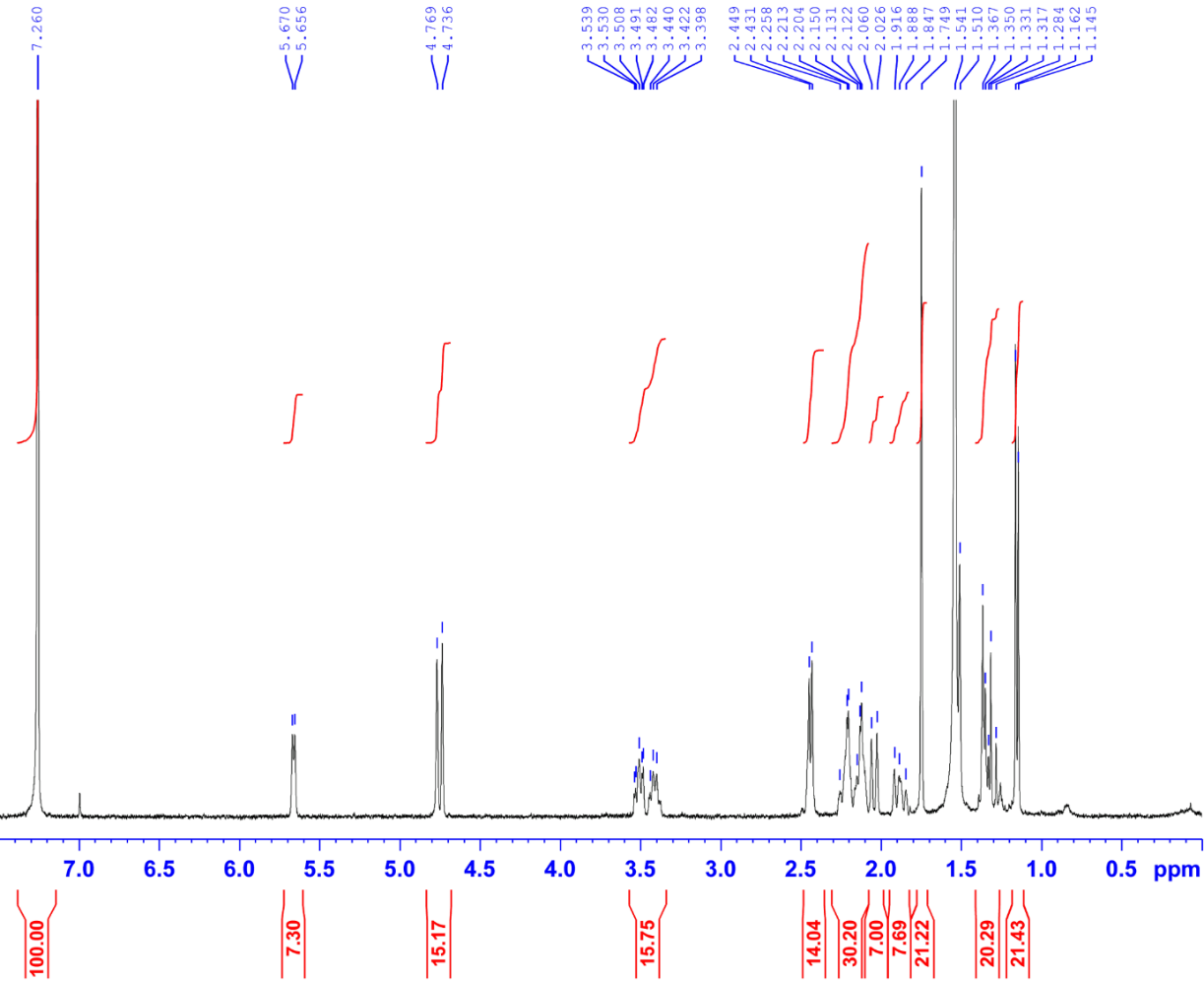

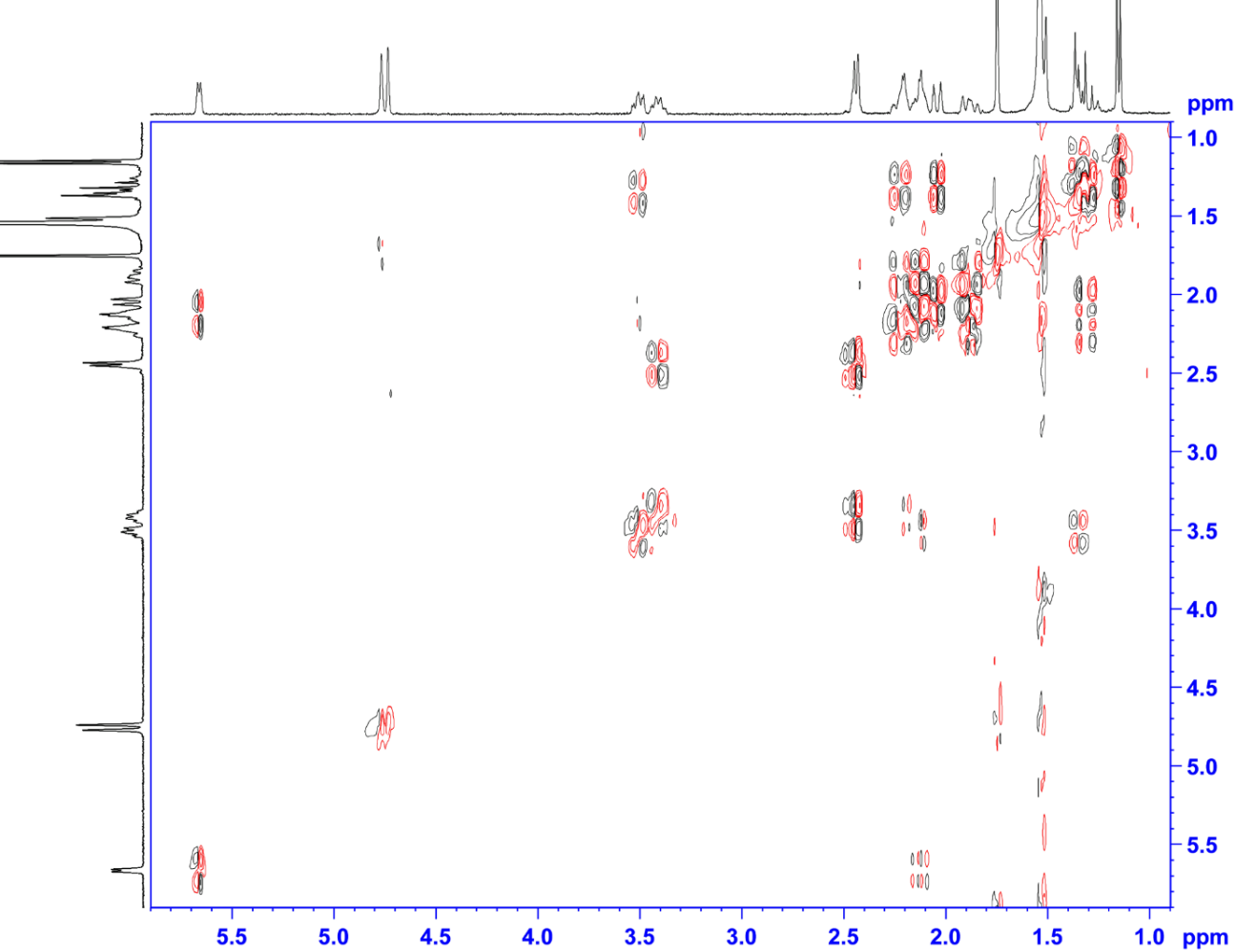

**Supplementary Fig. S8.** NMR of metabolite **5** (in CDCl_3_ at 400 MHz). (from top) ^1^H NMR, DQF-COSY.

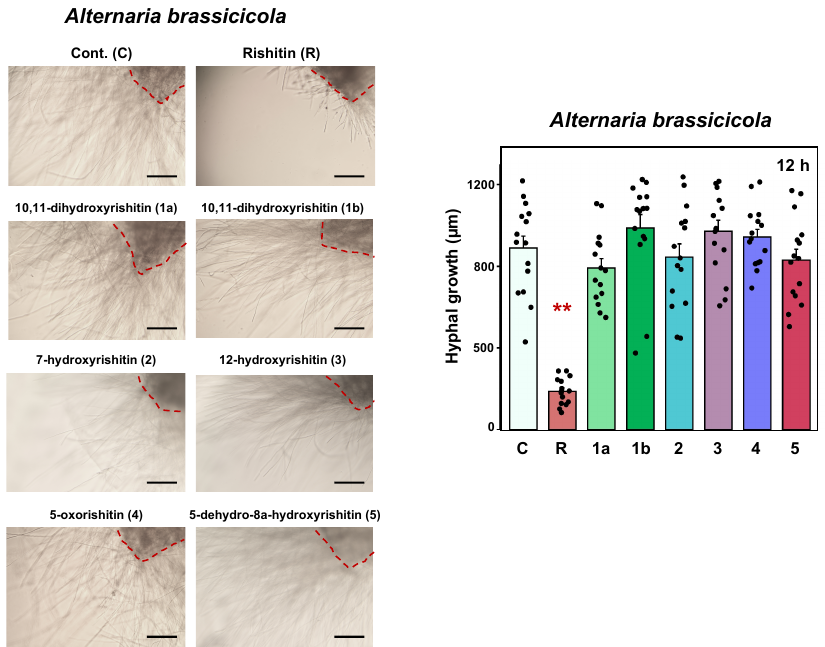

**Supplementary Fig. S9.** Anti-microbial activity of rishitin metabolites against phytopathogenic fungi *Alternaria brassicicola*. Mycelial blocks (approx. 1 mm^3^) of *A. brassicicola* were incubated in 50 µl of 0.1% DMSO (Cont.) or 500 µM rishitin or 500 µM rishitin metabolites. Outgrowth of hyphae from the mycelial block (outlined by dotted red lines) was measured after 12 h incubation. Bars = 200 µm. Data are means ± SE (n = 15). Data marked with asterisks are significantly different from the control as assessed by the two-tailed Student’s *t*-test. **P < 0.01.
